# Spatial inference of ancestor locations suggests northern refugia for canopy-forming kelps in the Northeast Pacific

**DOI:** 10.64898/2026.01.09.698529

**Authors:** Jordan B. Bemmels, Kristy J. Kroeker, Stephen R. Palumbi, Rachael A. Bay, Kristen M. Gruenthal, Sandra C. Lindstrom, Matthew M. Osmond, Gregory L. Owens

**Author notes:** Corresponding authors: JBB,; GLO.

## Abstract

- Pockets of the formerly glaciated Pacific coastline of North America likely remained ice-free throughout the Last Glacial Maximum (LGM). These areas may have served as refugia for terrestrial species, but less is known about their role in the persistence of coastal marine species.
- We examined genetic diversity from >1,000 newly and previously sequenced whole genomes of canopy-forming kelps of the genera *Nereocystis* and *Macrocystis*, and built simple ecological niche models. We then reconstructed ancestral recombination graphs and modelled the geographic locations of genetic ancestors through time.
- We detected high genetic diversity in both species in north-central British Columbia, in a region where suitable LGM habitat is plausible. Ancestor locations spatially converged backward in time toward this region, with multiple refugia inferred between northern Vancouver Island and southern Haida Gwaii. An expanded set of global samples for *Macrocystis* confirmed pre-LGM divergence with California but hinted at the possibility of subsequent gene flow.
- Kelp forests survived in northern refugia and have continuously occupied portions of the northern Pacific coastline since the LGM. One implication of these results is their support for the Kelp Highway Hypothesis of the peopling of the Americas by a coastal route.

## Introduction

Range shifts associated with Pleistocene glacial cycles have profoundly impacted species distributions and geographic patterns of genetic diversity (Hewitt, 2000; Soltis *et al*., 2006). Characterizing range shifts has been critical to explaining the evolution of genetically distinct lineages (Pamilo & Savolainen, 2004), understanding community assembly (Cortés-Guzmán *et al*., 2024), and prioritizing areas of elevated and endemic genetic diversity for conservation (Hampe & Petit, 2005; Médail & Diadema, 2009). Early studies of temperate terrestrial species emphasized a general pattern of range contraction during glacial periods into low-latitude refugia, and postglacial range expansion to higher latitudes as ice sheets receded (Davis, 1983; Hewitt, 2000). Founder effects during range expansion are expected to cause decreasing genetic diversity along the path of migration (Hewitt, 2000), but elevated diversity has also been observed in areas of secondary contact between genetically-differentiated lineages originating from separate refugia (Petit *et al*., 2003).

In recent decades, traditional expectations of southern macrorefugia for temperate Northern Hemisphere species have been challenged by the inferred existence of northern microrefugia located in unique microclimates close to glacial margins (Stewart & Lister, 2001; Provan & Bennett, 2008; Lumibao *et al*., 2017; Hošek *et al*., 2024). Terrestrial paradigms have also been extended to coastal marine species (Maggs *et al*., 2008; Neiva *et al*., 2016), where support has been found for both low- (Assis *et al*., 2018) and high-latitude refugia (Neiva *et al*., 2020; Bringloe *et al*., 2022; Jaugeon *et al*., 2025). Nonetheless, the refugial histories of many regions remain subject to debate. One such region (Mann & Gaglioti, 2024) is the formerly glaciated Pacific coastline of North America (Supporting Information Fig. S1). The Last Glacial Maximum (LGM) occurred asynchronously along different regions of this coastline, with local maxima reached from 24-15 ka (Mann & Hamilton, 1995; Mann & Gaglioti, 2024). During local maxima, terrestrial ice sheets largely extended up to or beyond the coastline (Mann & Hamilton, 1995), but several coastal pockets may have remained continuously unglaciated (Mann & Gaglioti, 2024). Geological and fossil evidence has suggested small unglaciated regions in the southern Alexander Archipelago (Carrara *et al*., 2007), Haida Gwaii (Warner *et al*., 1982; Mathewes & Clague, 2017), the central coast of British Columbia (BC) (Shaw *et al*., 2020), and northern Vancouver Island (Hebda *et al*., 2022), though these interpretations are not without controversy (Walcott *et al*., 2022; Mann & Gaglioti, 2024).

Genetic evidence has sometimes supported the idea that these putatively unglaciated pockets along the Pacific coastline served as refugia for terrestrial (Shafer *et al*., 2010) and marine (Jacobs *et al*., 2004; Marko *et al*., 2010) species. Terrestrial refugia in the southern Alexander Archipelago and Haida Gwaii may have hosted diverse taxa (Shafer *et al*., 2010) including large and small mammals (Heaton *et al*., 1996; Byun *et al*., 1997; Sawyer *et al*., 2019; Colella *et al*., 2021), birds (Pruett *et al*., 2013; Geraldes *et al*., 2019), freshwater fish (McCusker *et al*., 2000), and various angiosperms and ferns (Soltis *et al*., 1997). Marine species that may have survived off adjacent coastlines include intertidal fish and invertebrates (Hickerson & Ross, 2001; Hickerson & Cunningham, 2005; Marko *et al*., 2010), coho salmon (Smith *et al*., 2001), and red and brown algae (Lindstrom *et al*., 1997; Gierke *et al*., 2023). Additional marine refugia may have been located in the Gulf of Alaska (Grant *et al*., 2020; Grant & Bringloe, 2020; Grant & Chenoweth, 2021) and the central coast of BC (Lindstrom *et al*., 2021, 2025). However, relative to terrestrial taxa from this region, marine species have been less frequently studied and their phylogeographic histories are often contradictory (Marko *et al*., 2010). In addition, most existing marine studies from this region have relied on qualitative interpretation of genetic diversity patterns from a few loci, and even recent studies have typically not adopted a genomic perspective (Grant *et al*., 2020; Grant & Bringloe, 2020; Grant & Chenoweth, 2021; Gierke *et al*., 2023).

Genomic studies would be well suited to address outstanding controversies (Walcott *et al*., 2022; Mann & Gaglioti, 2024) about the potential importance of northern refugia to the phylogeographic history of the Northeast Pacific. Ideally, genomic studies would leverage recent conceptual and statistical advances that allow characterization of past species distributions and migration histories in previously unprecedented detail. Such advances have included, for example, the development of the directionality index (*ψ*) to infer the direction of range expansion (Peter & Slatkin, 2013, 2015); the integration of demographic and genetic models to infer the origins of range expansions and other parameters in a spatial landscape (He *et al*., 2017; Becheler & Knowles, 2022); and the emerging use of ancestral recombination graphs (ARGs) to reconstruct the recombination and coalescence history of all loci across the genome (Lewanski *et al*., 2024) and then infer the geographic locations of genetic ancestors through time (Wohns *et al*., 2022; Osmond & Coop, 2024; Deraje *et al*., 2025; Grundler *et al*., 2025). In addition, studies that focus on widespread, ecologically important taxa as representatives of coastal marine ecosystems would be particularly valuable.

Kelp forests represent one of the major temperate marine ecosystems of the Northeast Pacific. The two canopy-forming species from this region are bull kelp (*Nereocystis luetkeana*), distributed from central California to southwest Alaska, and giant kelp (*Macrocystis* spp.), distributed from Baja California to southern Alaska as well as in the Southern Hemisphere (Macaya & Zuccarello, 2010). Like other kelps, these foundation species create habitat for highly biodiverse communities (Teagle *et al*., 2017) and promote ecosystem function through nutrient cycling and high primary productivity (Wernberg *et al*., 2019; Eger *et al*., 2023). Both species have been culturally and economically important to humans for many millennia (Turner, 2001; Dillehay *et al*., 2008), with the Kelp Highway Hypothesis proposing that kelp forests along the margins of deglaciating landscapes (after ∼17 ka) could have provided vital resources to support initial human migration into North America (Erlandson *et al*., 2007; Braje *et al*., 2017).

Previous research on *Nereocystis* and *Macrocystis* has suggested divergent phylogeographic histories. A Haida Gwaii or Southeast Alaska refugium was inferred for *Nereocystis* based on high genetic diversity of seven microsatellite markers (Gierke *et al*., 2023), though diversity was fairly high along many regions of the outer coast. In contrast, Assis *et al*. (2023) considered all *Macrocystis* populations north of Oregon to be of postglacial origin based on low genetic diversity among six microsatellite loci and a global ecological niche model. However, few northern populations were sampled and genetic diversity was higher in Alaska than BC, hinting that the postglacial history of *Macrocystis* may be more complicated than range expansion from a single southern refugium.

The relationships among *Macrocystis* populations are also complicated by the presence of four holdfast morphologies. Of the two morphs present in North America, the *pyrifera* morph occurs from Baja to central California, and disjunctly in portions of Alaska (Gonzalez *et al*., 2023) and Haida Gwaii (Saunders & McDevit, 2014); the *integrifolia* morph occurs from central California to Alaska (Macaya & Zuccarello, 2010). Both morphs also occur in the Southern Hemisphere. Whether morphs represent ecotypes of a monotypic global species *M. pyrifera* (Demes *et al*., 2009) or multiple species (Lindstrom, 2023) remains controversial. It was recently demonstrated that *integrifolia* and *pyrifera* morphs from the same locality in California are genetically distinguishable (Gonzalez *et al*., 2023) and that holdfast morphology is genetically determined (Gonzalez & Raimondi, 2024). Current taxonomy considers all *Macrocystis* north of Point Conception, California to be *M. tenuifolia* but recognizes that genetic relationships within the Northern Hemisphere remain unclear (Lindstrom, 2023).

To reconstruct the history of canopy-forming kelp forests in the formerly glaciated Northeast Pacific, we analyzed newly and previously sequenced whole genomes of *Nereocystis* and *Macrocystis* from Alaska, British Columbia, and Washington (hereafter, AKBCWA) and globally for *Macrocystis.* We characterized patterns of genetic diversity, divergence, and gene flow among populations and constructed ARGs to infer population demographic history and the locations of genetic ancestors. Our main aims were 1) to infer whether refugia were located only in southern areas or also in northern areas along glacial margins, 2) to identify specific refugial locations, and 3) to test whether AKBCWA and California *Macrocystis* populations of either morph have been in recent genetic contact by estimating divergence times and testing for gene flow. Overall, these analyses will reveal how long canopy-forming kelp forests have been present along formerly glaciated northern coastlines, including in relation to proposed timelines for human migration into North America.

## Materials and methods

### DNA sampling and sequencing

We analyzed 505 *Nereocystis* and 522 *Macrocystis* whole-genome sequences (Supporting Information Tables S1, S2) from previously published datasets and de novo sequencing. We newly sequenced 56 *Nereocystis* and 21 *Macrocystis* from Alaska using previously described protocols (Bemmels *et al*., 2025). Briefly, we extracted DNA from silica-dried blade tissue using a custom CTAB method (Bemmels *et al*., 2025) and prepared libraries for *Illumina* sequencing using *Integrated DNA Technologies (IDT)* xGen DNA Library Prep EZ Kits. Paired-end 150-bp libraries were sequenced on an *Illumina* NovaSeq X at Canada’s Michael Smith Genome Sciences Centre (Vancouver). In addition, we performed low-depth whole-genome sequencing (for further details see Supporting Information Methods S1) of 137 new *Macrocystis* samples from California (Supporting Information Table S1), with libraries prepared following Rowan *et al*. (2019).

The North American *Macrocystis* samples represented in this study are a mixture of verified *integrifolia,* verified *pyrifera,* and unknown morphs (Supporting Information, Tables S1, S2). In California, all previously sequenced populations we analyzed are *pyrifera* morphs, except for a single *integrifolia* population from Stillwater Cove (Gonzalez *et al*., 2023). Morph identity was not assessed for any of the newly sequenced California populations. Morph identity was also not assessed for any previously sequenced BC and Washington populations, but these individuals are presumably mostly *integrifolia* as this is the primary morph from this region, except in Haida Gwaii where both morphs are known (Saunders & McDevit, 2014). In contrast, all newly sequenced Alaska populations are *pyrifera* morphs (S.C. Lindstrom, personal communication).

### SNP genotyping

We generated single nucleotide polymorphism (SNP) genotypes following Bemmels et al. (2025). We removed adapter contamination and filtered raw reads using default parameters in *fastp* v.0.23.2 (Chen *et al*., 2018) and aligned reads to reference genomes for *Nereocystis* (NCBI: GCA_031213475.1) (Alves-Lima *et al*., 2025) and *Macrocystis* (JGI PhycoCosm: *Macrocystis pyrifera* CI_03 v1.0) (Diesel *et al*., 2023) using default parameters in *bwa-mem* 0.7.18-r1243 (Li, 2013). We merged all aligned reads for each individual and removed duplicates using *SAMtools* v.1.22.1 (Danecek *et al*., 2021) and *Picard* v.2.26.3 (Broad Institute, 2024), respectively. To correct for read misalignments around indels, we realigned reads using the *RealignerTargetCreator* and *IndelRealigner* tools from GATK v.3.8 (Van der Auwera & O’Connor, 2020).

We called genotypes using *BCFTools* v.1.17-1.22 (Danecek *et al*., 2021) with the commands *bcftools mpileup -Q 30 -q 30* and *bcftools call -m -v,* and filtered calls to biallelic SNPs with the command *bcftools view -v snps -m 2 -M 2 -q 0.000001:minor*. To remove sites that likely represent alignment errors and confounded loci, we (1) used *BCFTools* to remove sites with >50% heterozygosity; (2) retained only SNPs passing additional quality metrics (SGB ≥ 2; −1.96 ≤ x ≤ 1.96, where x is each of MQBZ, MQSBZ, and RPBZ, respectively; and VDB ≥ 0.05), following Bemmels *et al*. (2025); and (3) removed sites above the 98^th^ percentile of overall depth.

To this set of quality-filtered sites, we subsequently applied missing-data filters. We created datasets retaining individuals with mean depth ≥8x for both species. For *Macrocystis* we additionally created a dataset with mean depth ≥1x, in order to include the low-depth samples from California. We then set genotypes to missing for all sites below 8x or 1x coverage, depending on the dataset, using the *bcftools +setGT* plugin, and removed all sites with ≥20% missing data. We hereafter refer to these datasets as the 8x and 1x datasets, respectively. For all datasets, we retained only autosomal scaffolds and contigs ≥1.5 Mbp in length.

We then identified and removed genetically identical individuals and close relatives using *ngsRelate* v.2 (Hanghøj *et al*., 2019). We ran *ngsRelate* on individual VCF files for each population and examined the pairwise relatedness *r*_ab_ (Hedrick & Lacy, 2015; Hanghøj *et al*., 2019). We considered close relatives to be those where *r*_ab_ > 0.3536. Although our study species frequently inbreed (Bemmels *et al*., 2025), in the absence of inbreeding, *r*_ab_ approximates twice the coefficient of kinship (Hedrick & Lacy, 2015). Thus, the threshold 0.3536 distinguishes first-degree from second-degree relatives (Manichaikul *et al*., 2010) with expected *r*_ab_ = 0.5 and 0.25, respectively. For clusters of related individuals, we retained only the individual with the smallest proportion of missing genotypes. Our final standard dataset sizes were as follows: 8x *Nereocystis*: 470 individuals, 16,628,760 SNPs; 8x *Macrocystis*: 311 individuals, 11,091,173 SNPs; 1x *Macrocystis*: 484 individuals, 12,639,657 SNPs. We also created datasets of invariant sites using the same methods as above, except that we removed the -v flag from *bcftools call*, retained only sites with one unique allele using *bcftools view -Q 0.000001:nonmajor -V indels*, and did not apply any filters that are applicable to polymorphic sites only.

Some downstream analyses required removing selfed individuals, assuming that selfing reflects dispersal limitations rather than non-random mating in kelp (Gaylord *et al*., 2006; Edwards, 2022). We identified selfed individuals from runs of homozygosity (ROHs) using *bcftools roh.* We used the 8x datasets, with repetitive regions of the genome previously identified by Bemmels *et al*. (2025) removed from input VCFs using the *intersect* command in *BEDTools* v.2.30.0 (Quinlan & Hall, 2010). We then calculated the inbreeding coefficient *F*_ROH_ as the proportion of the genome in ROHs and considered individuals with *F*_ROH_ > 0.3536 to be selfed (Manichaikul *et al*., 2010) but reclassified any individuals with *F*_ROH_ > 0.3536 as non-selfed if homozygous regions were dispersed in small segments rather than large contiguous stretches based on visual inspection of observed heterozygosity across the genome.

### Population genetic structure

For AKBCWA, we performed principal component analysis (PCA) using *EMU* v. 1.2.1 (Meisner *et al*., 2021). We used the 8x datasets with SNPs filtered to a minimum minor allele frequency (MAF) of 0.01 and thinned to a minimum distance of 10 kbp, prior to running *EMU* using 1000 iterations (*--iter 1000*) and three eigenvectors for missing data estimation (*-e 3*). For *Macrocystis* analyses including low-depth California samples, we used the read-sampling PCA approach implemented in *ANGSD* v.0.941 (Korneliussen *et al*., 2014) with BAM input but the same set of sites (*-sites* flag) previously identified in the 1x dataset. We generated a covariance matrix in ANGSD using the flags *-minMapQ 30 -minQ 30 -doMajorMinor 1 -doMaf 1 -doIBS 1 -doCov 1 - doCounts 1 -makeMatrix 1 -GL 1*. We then calculated eigenvalues and eigenvectors from the covariance matrix using the *eigen()* function in *R* v.4.4.2 (R Core Team, 2024).

We identified genetic clusters in AKBCWA using *fastSTRUCTURE* v.1.0 (Raj *et al*., 2014). We used the 8x datasets and performed an initial 100 runs for each number of clusters (*K*) from *K*=1 to 10. We selected the run with the highest likelihood for each *K*-value and selected the optimal value using the script *chooseK.py* (distributed with *fastSTRUCTURE*). For both species, the optimal *K*-value was 8. However, we selected *K*=7 as our final result because the eighth cluster represented a locally bottlenecked population rather than a larger regional grouping in *Nereocystis*, and was geographically discontiguous and possibly a statistical artifact in *Macrocystis*.

We calculated population nucleotide diversity (*π*) from the global 8x datasets including invariant sites for each species using *pixy* v.1.2.7.beta1 (Korunes & Samuk, 2021). We also used *pixy* to calculate pairwise genetic divergence (*d*_XY_) (Nei & Li, 1979) between genetic clusters. We calculated pairwise genetic differentiation (*F*_ST_) (Weir & Cockerham, 1984) using *hierfstat* v.0.5-11 (Goudet, 2005), from the 8x datasets with minimum MAF 0.01 and thinned to 10 kbp. We also calculated the directionality index (*ψ*) (Peter & Slatkin, 2013) in AKBCWA using the 8x dataset for each species, filtered to minimum MAF of 0.05 and thinned to minimum 10 kbp. ψ infers the direction of range expansions between populations, given shifts in allele frequencies that occur during founder events. We polarized alleles for *ψ* calculation following Bemmels *et al*. (2025) from alignments to outgroup genomes (Grigoriev *et al*., 2021; Phaeoexplorer Project, 2024), using *bwa-mem* v.0.7.17-r1188 (Li, 2013), *SAMtools* v.1.17 (Danecek *et al*., 2021), *HTSBox* v.r345 (Li, 2012), and scripts from Taylor *et al*. (2024). For further details see Supporting Information, Methods S1.

To further explore the relationship between *Macrocystis* from AKBCWA and California, we constructed a phylogeny (Lewis, 2001; Kalyaanamoorthy *et al*., 2017) and assessed branch support (Guindon *et al*., 2010; Hoang *et al*., 2018) using *IQ-TREE* v.2.3.6 (Minh *et al*., 2020) from SNPs thinned to 10-kbp retaining one individual per population. We tested for the possibility of gene flow between different geographic regions using four-taxon Patterson’s *D*-statistics (Durand *et al*., 2011) inferred in *Dsuite* v.0.5 r53 (Malinsky *et al*., 2021). For further details see Supporting Information, Methods S1.

### Ecological niche modelling

To explore where climatically suitable habitat may have existed for kelp during the LGM, we constructed simple ecological niche models (ENMs) using *MaxNet* v.0.1.4 (Phillips *et al*., 2006, 2017) within the *R* package *SDMtune* v.1.3.1 (Vignali *et al*., 2020). We used occurrence records for both species from the Global Biodiversity Information Facility and controlled for spatial bias using brown algae (Phaeophyceae) as background points (GBIF.org, 2024a,b,c). We used environmental predictor variables characterizing temperature and salinity regimes from MARSPEC (Braconnot *et al*., 2007; Sbrocco & Barber, 2013; Sbrocco, 2014). For further details see Supporting Information, Methods S1.

### Ancestral recombination graphs

We constructed ancestral recombination graphs (ARGs) (Lewanski *et al*., 2024) using *Relate* v.1.2.3 (Speidel *et al*., 2019, 2021). Full details of ARG construction are provided in Supporting Information, Methods S1. In brief, we used the 8x datasets filtered to AKBCWA (with SNPs polarized as described in the *Pairwise directionality index details* section of Supporting Information, Methods S1), removed selfed individuals (Mather *et al*., 2020), phased and imputed missing data using *Beagle* v.5.5 (Browning *et al*., 2018, 2021), and estimated the required recombination map using *FastEPRR* v.2.0 (Gao *et al*., 2016). We placed an initial prior on the mutation rate of 8.135 x 10^-10^ for both species, following Bemmels *et al*. (2025), and priors on haploid effective population size of 2N = 200,000 and 20,000 for *Nereocystis* and *Macrocystis*, respectively. For rescaling results to absolute time in years, we assumed a generation time of 1 year in the annual species *Nereocystis* and 2 years in the perennial species *Macrocystis* based on qualitative interpretation of life history and ecological data (Dayton et al., 1984; Reed, 1987; Bell & Siegel, 2022), as outlined in Supporting Information, Methods S1. We created initial ARG topologies in *Relate* (*--mode all*) and then simultaneously re-estimated branch lengths, effective population sizes (*N*_e_), and mutation rates using the *Relate* script *EstimatePopulationSize.sh*. We also repeated ARG reconstruction for *Macrocystis* using the global 8x dataset thinned to one population per genetic cluster from AKBCWA and all populations from Chile and California.

We estimated the demographic history of each population and the timing of population splits using the output from *EstimatePopulationSize.sh*. We calculated *N*_e_ in each time bin as 0.5/*r*, where *r* is the haploid coalescence rate (Speidel *et al*., 2019), and took the mean across all chromosomes. We also calculated the relative cross-coalescence rate (rCCR) as the coalescence rate between populations divided by the mean coalescence rate within populations (Schiffels & Wang, 2020), but arbitrarily set the upper limit on rCCR for each chromosome to two prior to averaging across chromosomes, to prevent rare extreme outliers (that arise when comparing ratios of very small numbers) from distorting overall patterns. We estimated divergence time between populations as the midpoint of the most recent time bin at which the rCCR first rose above 0.5 (Schiffels & Wang, 2020).

### Inferred ancestral locations

We used *Spacetrees* (Osmond & Coop, 2024) to infer the geographic locations of genetic ancestors through time from the ARGs for AKBCWA. *Spacetrees* spatially models Brownian motion of ancestors down trees in an ARG. To reduce correlations between trees we first thinned to every 500^th^ tree, resulting in 844 trees for *Nereocystis*, and to every 100^th^ tree, resulting in 887 trees for *Macrocystis*. To account for uncertainty in branch length estimates, each of these trees was sampled 100 times (using *Relate*’s *SampleBranchLengths.sh* script). Ancestor locations were then inferred using *Spacetrees*’ best linear unbiased predictor (BLUP) method, which averages the most likely location over samples of each tree, as this method is fast and robust to outlier branch-length estimates. For visualization, we calculated mean ancestor locations across trees for each individual in the entire dataset (all AKBCWA samples) and for each genetic cluster. For each genetic cluster, we also calculated spatial contour lines containing 90% of all inferred ancestor locations using the *contour()* function in *R* v.4.4.2 (R Core Team, 2024) from a kernel density of 100×100 pixels generated with the *kde2d()* function in the *R* package *MASS* (Venables & Ripley, 2002).

## Results

### Population genetic structure

Genetic clusters identified in *fastSTRUCTURE* and PCAs of genetic variation corresponded closely with geography and identified a close relationship between northern BC and newly sequenced Alaska samples in both species (Fig. 1). Among North American *Macrocystis*, the first principal component (PC) axis of variation separated AKBCWA from verified *pyrifera* and unassessed Californian individuals, with the single verified *integrifolia* individual from California falling in an intermediate position between AKBCWA and other Californian individuals (Fig. 2a,b). Within California, genetic variation among *pyrifera* and unassessed populations primarily followed a latitudinal gradient (Fig. 2c), except that the northernmost population from Bodega Bay (purple) unexpectedly clustered with the offshore Channel Islands (pink). Within AKBCWA, genetic diversity (*π*) was highest in north-central BC for both species (Fig. 3a,b). Globally, the highest *Macrocystis* diversity was found in California and was similar for both morphs (Fig. 3c), with intermediate diversity in AKBCWA and Chile and lowest diversity in Australia.

**Fig. 1.**
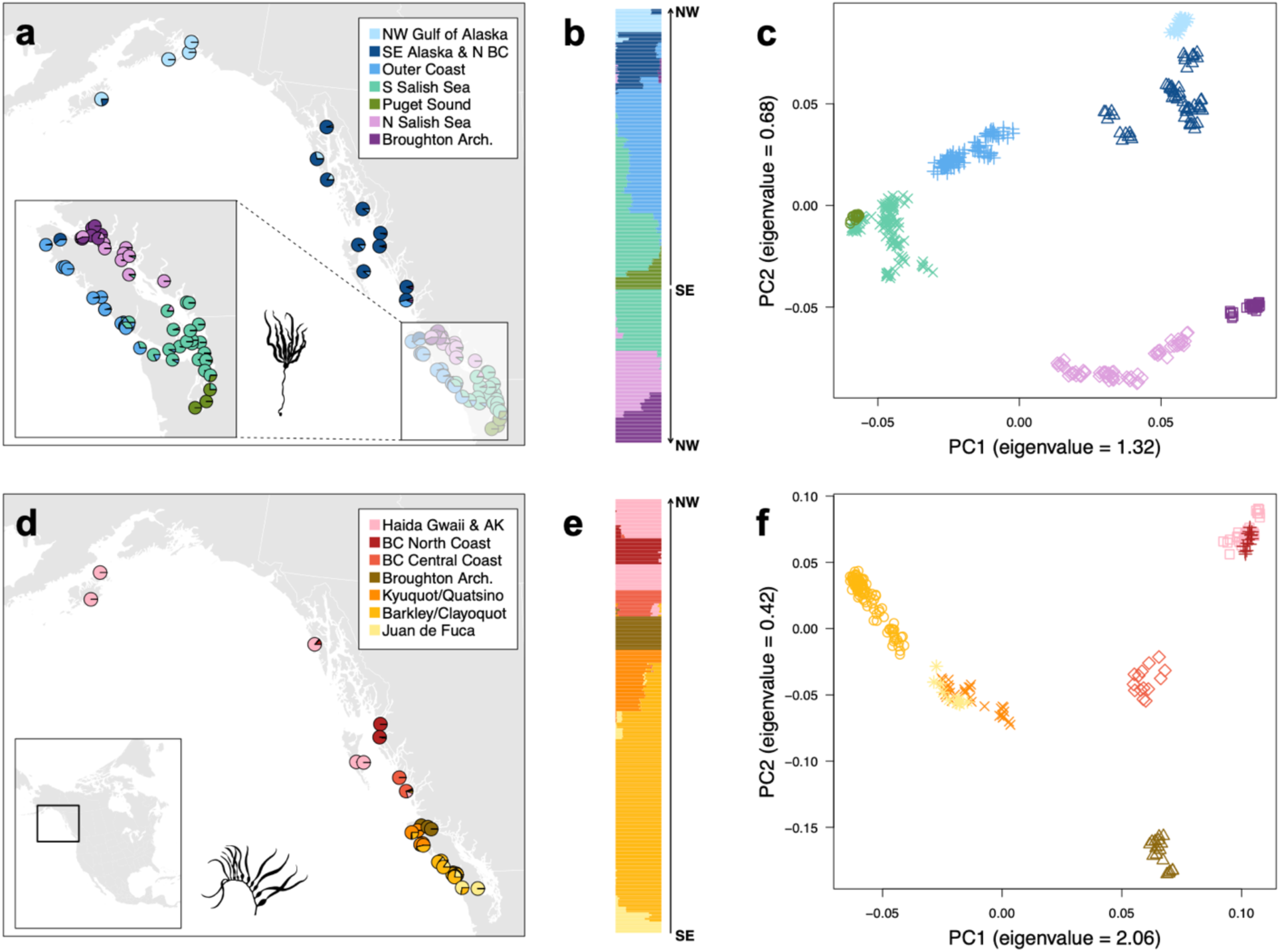
Population genetic structure in AKBCWA for (a-c) *Nereocystis* and (d-f) *Macrocystis*. (a,d) Genetic clusters identified in *fastSTRUCTURE*, with pie charts showing the proportion of each population’s ancestry in each genetic cluster. AK: Alaska; arch.: archipelago. (b,e) Ancestry in genetic clusters with each horizontal line representing one individual. Individuals are arranged geographically from southeast (SE) to northwest (NW). (c,f) PCA of genetic variation, with each point representing an individual and each genetic cluster represented by a different symbol and colour.

**Fig. 2.**
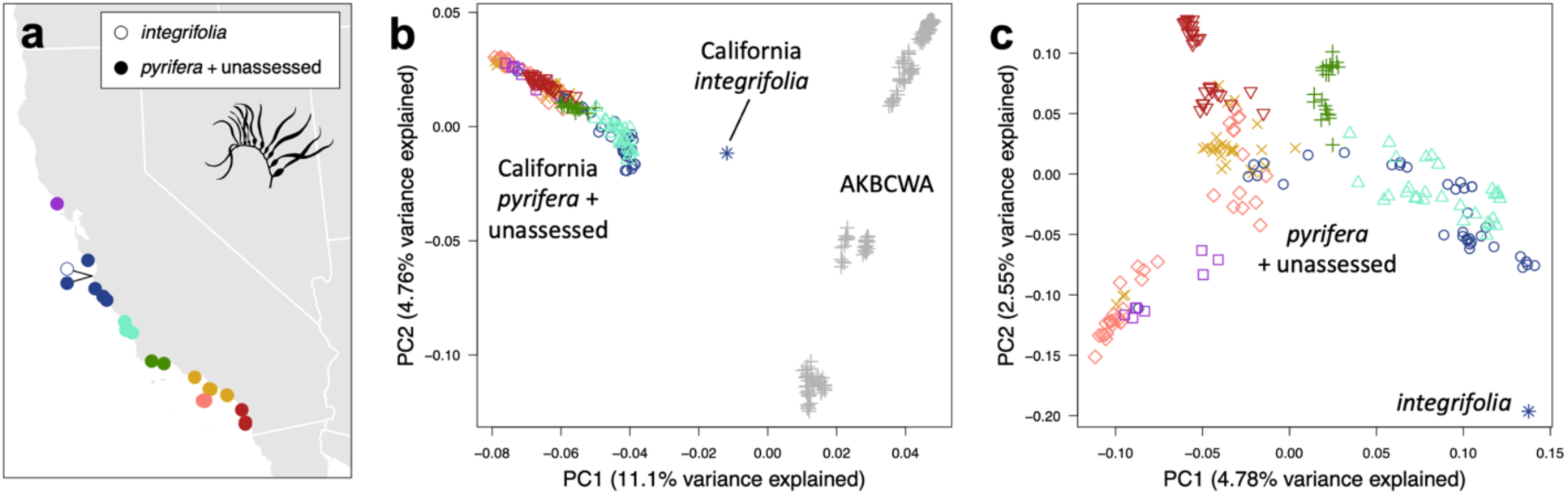
Population genetic structure of *Macrocystis* in California. (a) Map of populations showing colours used to represent individuals in (b) and (c). Open circle: *integrifolia* morph; closed circles: *pyrifera* morph and populations for which morph identity was unassessed. (b) PCA of genetic variation in California and AKBCWA. Note that AKBCWA includes both *pyrifera* (Alaska) and unassessed but presumed *integrifolia* (southern BC and Washington) morphs. (c) PCA of genetic variation in California only.

**Fig. 3.**
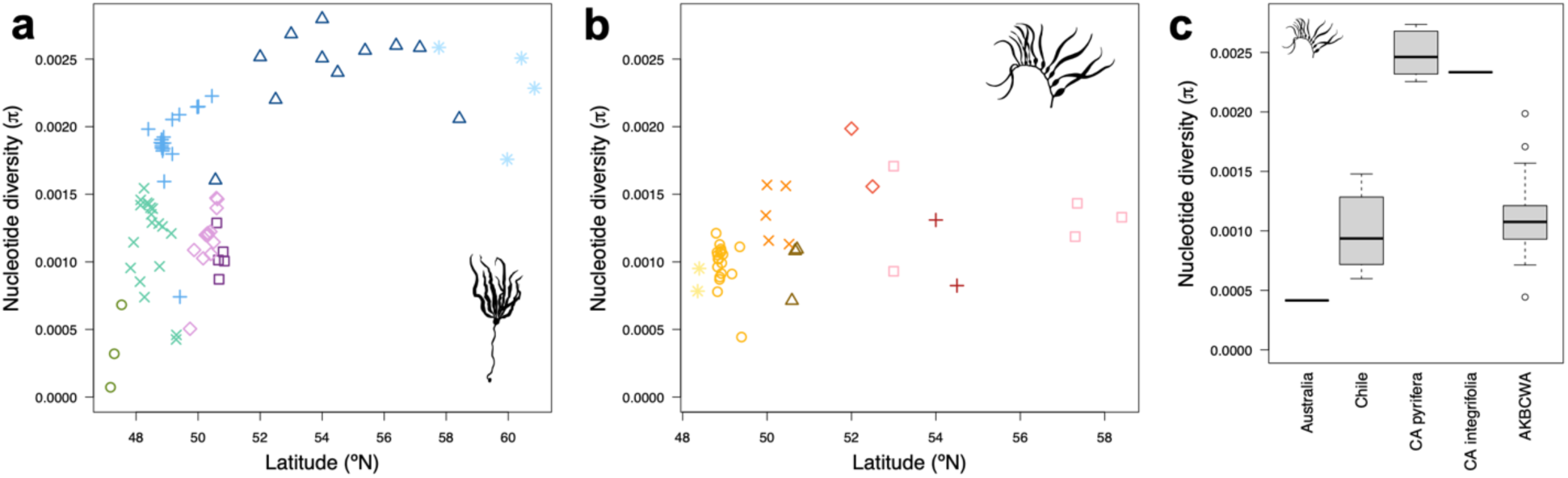
Geographic patterns of genetic diversity. (a,b) Relationship between population nucleotide diversity (*π*) and latitude (°N) in AKBCWA for (a) *Nereocystis* and (b) *Macrocystis*. Colours and symbols correspond to genetic clusters from Fig. 1. (c) Variation in π among *Macrocystis* populations from different global regions. CA: California.

Genetic divergence (*d*_XY_) among genetic clusters and regions largely corresponded to geography (Supporting Information Fig. S2), with more distant populations exhibiting higher *d*_XY_. Among global *Macrocystis*, *d*_XY_ was much higher between hemispheres than within hemispheres. The *d*_XY_ between southern AKBCWA individuals and California *integrifolia* was lower than between southern AKBCWA and California *pyrifera*, or between the two morphs within *California*. Genetic differentiation (*F*_ST_) did not show a consistent geographic pattern as it appeared strongly impacted by *π*, with low-diversity populations exhibiting high *F*_ST_ (Supporting Information Fig. S2).

Pairwise directionality indices (*ψ*) revealed similar inferred directions of postglacial range expansion in both species (Fig. 4), beginning in Haida Gwaii for *Nereocystis* and the central coast of BC for *Macrocystis* and proceeding in both a northerly and southerly direction. As expected for recent range expansions (Kemppainen *et al*., 2024), pairwise *ψ* was highly correlated with the difference in nucleotide diversity (Δ*π*) between populations (Fig. 4).

**Fig. 4.**
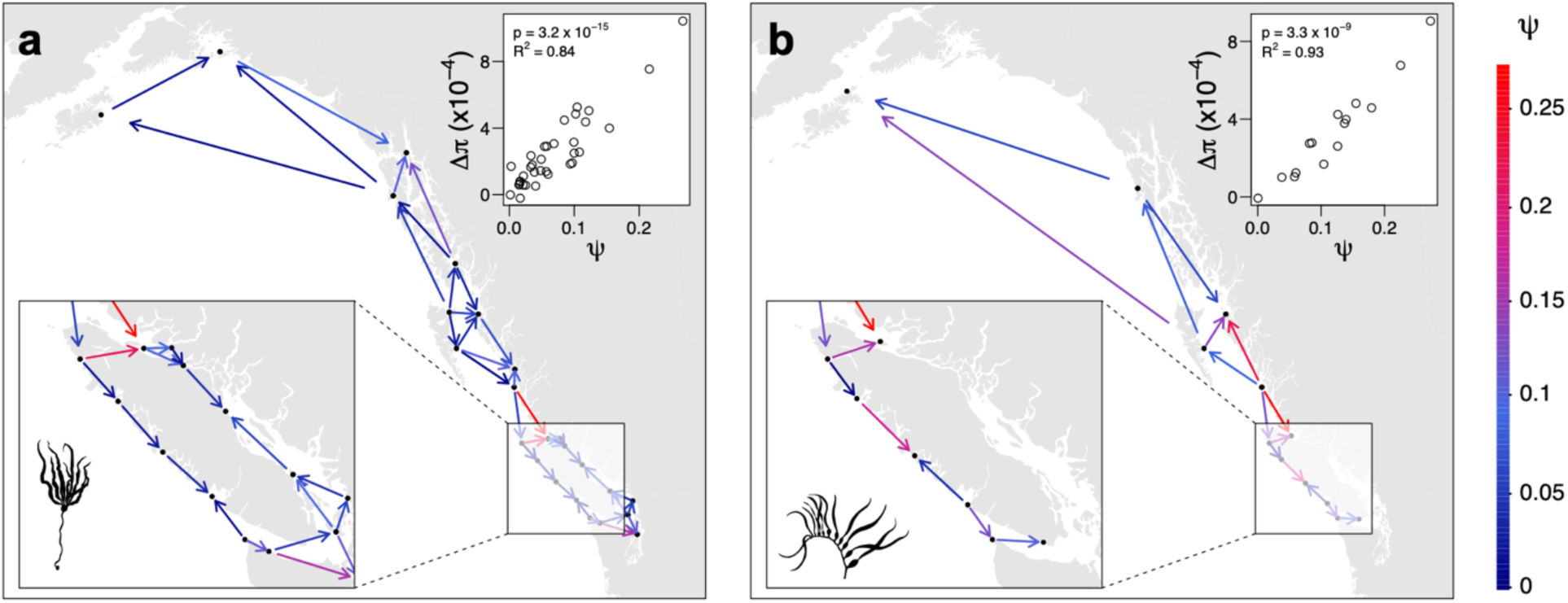
Directionality index (*ψ*) between pairs of populations of (a) *Nereocystis* and (b) *Macrocystis*. Arrows point in the inferred direction of range expansion and are coloured according to the magnitude of *ψ*, with higher *ψ* indicating a stronger signal of range expansion. Inset plots show the relationship between the population pairwise difference in nucleotide diversity (Δ*π*) and *ψ*.

### Macrocystis global phylogeny and gene flow

The *IQ-TREE* phylogeny of *Macrocystis* revealed a deep split between northern and southern hemispheres (Supporting Information Fig. S3a). Within North America, southern California *pyrifera* occupied the earliest diverging lineages. Northern California *pyrifera* was sister to a clade containing northern California *integrifolia* and AKBCWA. Support for clades that distinguished major global geographic regions was always high (SH-aLRT ≥ 80% and ultrafast boostrap support ≥ 95%), but branching patterns within smaller geographic regions were often poorly resolved (Supporting Information Fig. S3a).

Statistically significant signatures of gene flow were inferred in *Dsuite* between *integrifolia* and *pyrifera* morphs in northern California (Box 1; Supporting Information Fig. S3c) as well as between California *integrifolia* and southern AKBCWA populations (Box 2; Supporting Information Fig. S3c). Numerous other instances of gene flow were also inferred, mostly between geographically adjacent populations. Strong signals of gene flow between central BC (MP-CC-01 and MP-PH-01) and all individual southern BC populations may reflect the uncertain phylogenetic placement of central BC (Supporting Information Fig. S3a) or a non-tree-like relationship among populations from this region.

### Ecological niche models

Spatial biases in sampling effort were qualitatively similar between the focal species and brown algae that were used as background points to control this bias (Supporting Information Fig. S4). For both species, the same three final predictor variables were retained: mean annual sea surface salinity (SSS), sea surface temperature (SST) of the warmest ice-free month, and annual range in SST (Supporting Information Table S3). Predictive performance (Supporting Information Table S4) was moderate in terms of testing area under the receiver operating characteristic curve (AUC: 0.78-0.80; moderate AUC category: 0.7-0.9; Smith *et al*., 2021) and fair to moderate in terms of true skill statistic (TSS: 0.52-0.54; fair to moderate TSS category: 0.2-0.6; Smith *et al*., 2021).

The predicted distributions of both species in the current time period matched known distributions well (Supporting Information Figs. S4, S5), especially in *Nereocystis*. In *Macrocystis*, the model overpredicts presence (though with only marginal suitability) in some areas where it does not occur, including inner waterways (e.g., the inner Salish Sea and innermost Southeast Alaska) and west of Kodiak Island (Supporting Information Fig. S5b). The ENMs projected to the LGM ensemble model excluding CCSM3 (see Supporting Information, Methods S1) predicted marginally suitable habitat along the coastline of Alaska and BC in *Nereocystis* (Supporting Information Fig. S6a). In *Macrocystis*, the LGM ensemble projections predicted almost complete absence from this region (Supporting Information Fig. S6b). However, when projected to the CCSM3 climate model, the coastlines of Alaska and BC showed many areas of moderate habitat suitability in ice-free areas for *Nereocystis* and moderate to low suitability for *Macrocystis* (Supporting Information Fig. S7).

### Demographic and geographic inferences with ARGs

Effective population sizes (*N*_e_) inferred in *Relate* suggested late Pleistocene to Holocene bottlenecks in all populations of both species (Fig. 5a,b). In *Nereocystis*, bottlenecks typically reached minimum size near the Pleistocene-Holocene transition (11.7ka) (Fig. 5a). In *Macrocystis*, bottlenecks were more severe and more recent, with minima generally occurring between 10 and 1 ka. Divergence times among populations within AKBCWA were similar in both species (Fig. 5c,d). Median divergence times (Supporting Information Table S5) between population pairs within the same genetic cluster dated to the Holocene (*Nereocystis*: 3.2-8.6 ka; *Macrocystis*: 0.4-4.4 ka), while those between different genetic clusters were substantially older (*Nereocystis*: 6.1-119.5 ka; *Macrocystis*: 3.2-61.9 ka).

**Fig. 5.**
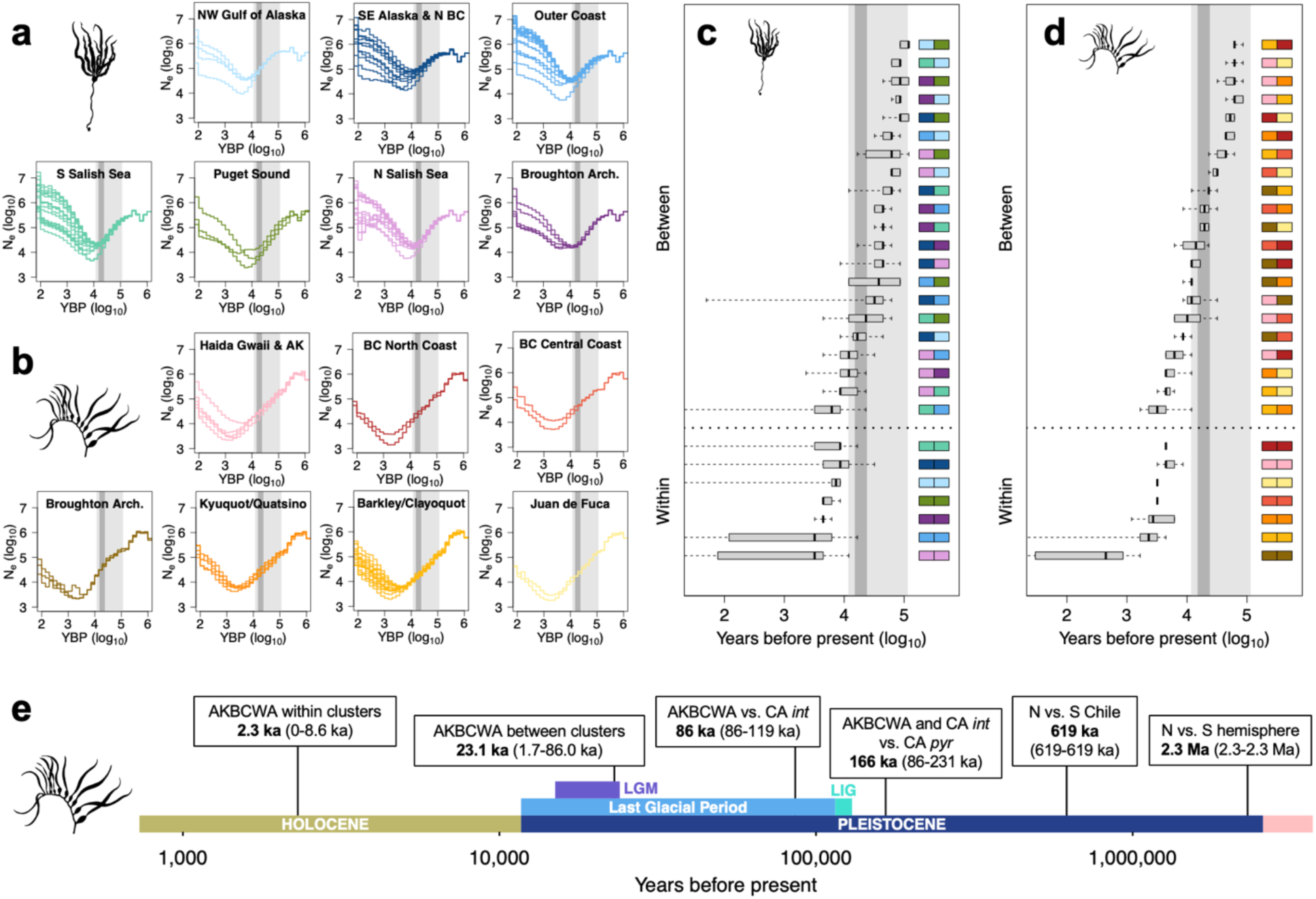
Population demographic events inferred from ARGs. (a,b) Changes in effective population size (*N*_e_) over time in populations of (a) *Nereocysits* and (b) *Macrocystis*, with populations grouped by genetic cluster. Light and dark grey horizontal bands show the timing of the Last Glacial Period and the local Last Glacial Maximum (LGM), respectively. AK: Alaska; arch.: archipelago; YBP: years before present. (c,d) Inferred population split times between pairs of populations within and between genetic clusters for (c) *Nereocystis* and (d) *Macrocystis* in AKBCWA. Pairs of genetic clusters are colour-coded as in (a) and shown on the right of the plot for each boxplot. Boxplots show the distribution of split times among all possible pairs of populations corresponding to each pair of genetic clusters, with whiskers extending to minimum and maximum values. See also Supporting Information Table S5 for a textual representation. (e) Timeline of inferred population split times between pairs of *Macrocystis* populations from different global regions, with median split times among all possible population pairs in bold and the corresponding range in parentheses. If the minimum and maximum of the range are identical, then all possible population pairs were inferred to have diverged in the same time bin. CA: California; int: *integrifolia* morph; pyr: *pyrifera* morph; LIG: Last Interglacial.

In *Macrocystis*, divergence times between populations from different global regions all dated to the Pleistocene (Fig. 5e). AKBCWA individuals and California *integrifolia* were estimated to have diverged at 86 ka (range among different population pairs: 86-119 ka), more recently than the divergence between AKBCWA and California *pyrifera* (166 ka; range: 119-231 ka) or between the two morphs within California (166 ka; range: 86-231 ka). Northern and southern Chilean populations diverged at 619 ka, while the Northern and Southern Hemisphere diverged at 2.3 Ma (with no range given for either estimate as the 50% rCCR corresponded to the same time bin for all pairs of populations).

The mean ancestor location inferred in *Spacetrees* toward which all populations collapse if allowed to coalesce toward completion (i.e., 10^6^ generations before present) is in southern Haida Gwaii for both species. This location should not be interpreted as the true mean ancestor location at 10^6^ generations because *Spacetrees* cannot accurately inference ancestor locations across multiple glacial cycles of range expansion and contraction (Osmond & Coop, 2024). However, the fact that it is substantially north of the mean latitude and longitude of sampled populations in the present day (northern Vancouver Island for both species) confirms that there is spatial signal in the ARGs. At shallower time scales, inferred locations of genetic ancestors for each genetic cluster showed both species increasingly converging toward central and northern BC backward in time (Fig. 6). The broad distribution of inferred ancestor locations at 20 ka for each genetic cluster highlights uncertainty about precise ancestor locations at the LGM. In addition, genetic coalescence may substantially predate spatial convergence of lineages, creating a lag in *Spacetrees* between arrival at a true ancestral population location and genetically inferred locations at that time. Thus, the generations prior to the LGM may help clarify precise refugial locations. In *Nereocystis*, mean ancestor locations from 20-40 ka for all but the northernmost genetic cluster converge toward two separate locations: northern Vancouver Island and southern Haida Gwaii (Fig. 6a). The northernmost cluster shows a separate far-northern location. In *Macrocystis* the mean ancestor locations from 20-80 ka converge toward three locations: southern Haida Gwaii, the central coast of BC, and northern Vancouver Island (Fig. 6b).

**Fig. 6.**
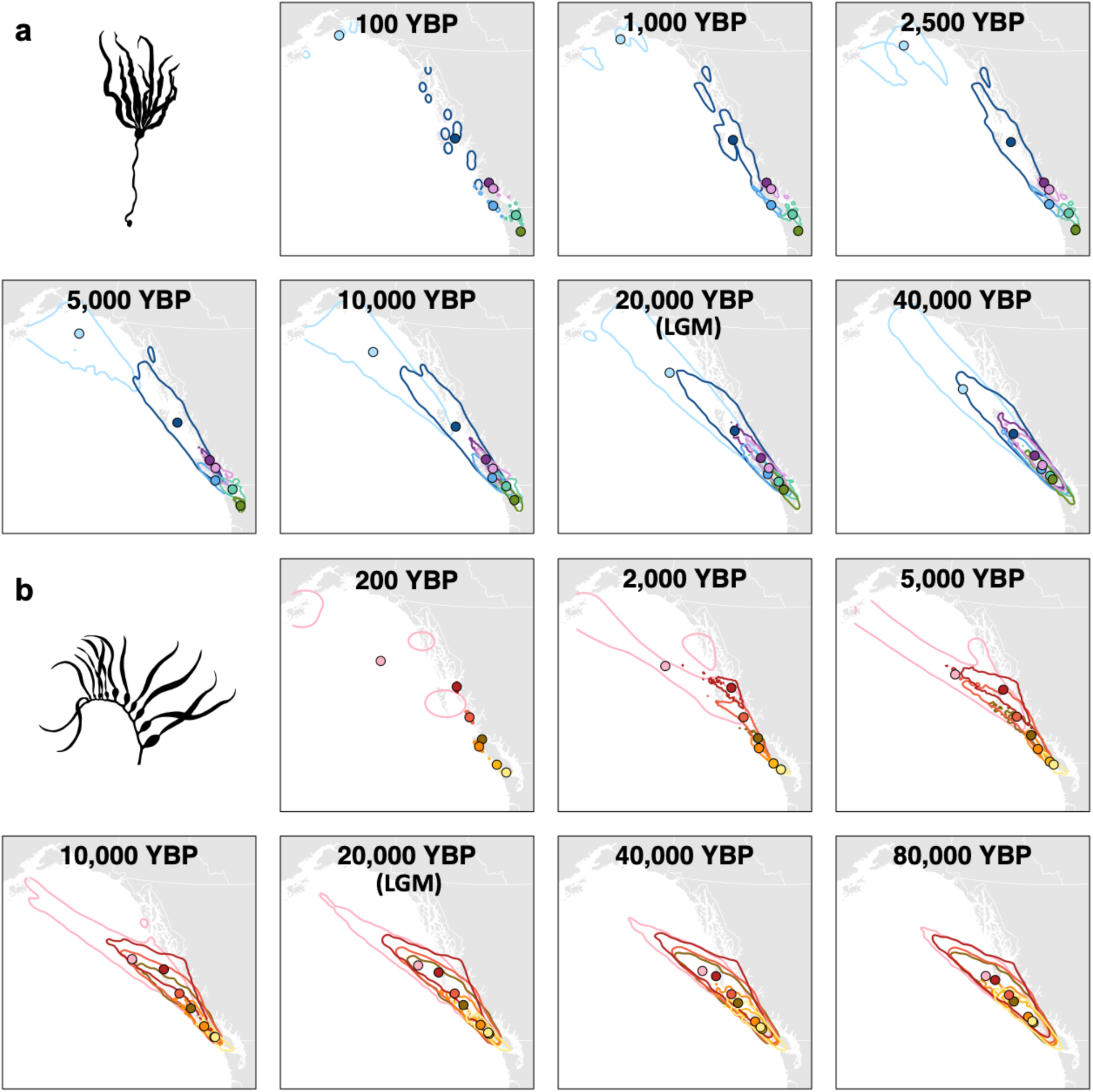
Geographic locations of ancestors through time inferred from ARGs using *Spacetrees* for (a) *Nereocystis* and (b) *Macrocystis*, visualized for each genetic cluster with colours corresponding to Fig. 1. Lines represent contours enclosing 90% of all ancestor locations for each genetic cluster, and circles represent the mean ancestor latitude and longitude. YBP years before present; LGM: Last Glacial Maximum.

## Discussion

Canopy-forming kelp forests likely persisted throughout the LGM in northern refugia along ice-free margins of the glaciated Pacific coastline of North America (Fig. 7). Range expansion from these refugia gave rise to contemporary populations of *Nereocystis* and *Macrocystis* from Washington to Alaska. ARG-based inference of ancestral locations using *Spacetrees* agreed with qualitative interpretations of genetic diversity, pairwise directionality indices, and simple ecological niche models, all of which highlighted the central to northern coast of BC as a likely refugial region. In addition, *Spacetrees* added increased resolution and suggested there may have been multiple refugia within this region, as ancestor locations of different populations converged toward either southern Haida Gwaii, the central coast of BC, or northern Vancouver Island. These regions were previously inferred from geologic evidence to have been continuously ice-free. Our results highlight the power of ARG-based models to geographically locate ancestors through time and reconstruct the origins of range expansions. Despite the inference of northern refugia, hints of gene flow in *Macrocystis* between southern AKBCWA and California (and between morphs within California) require further study. Overall, the presence of northern refugia for canopy-forming kelps suggests that kelp forests could potentially have supported rich coastal ecosystems throughout the LGM, which supports the hypothesis that these ecosystems may have been integral to the peopling of the Americas via a coastal route.

**Fig. 7.**
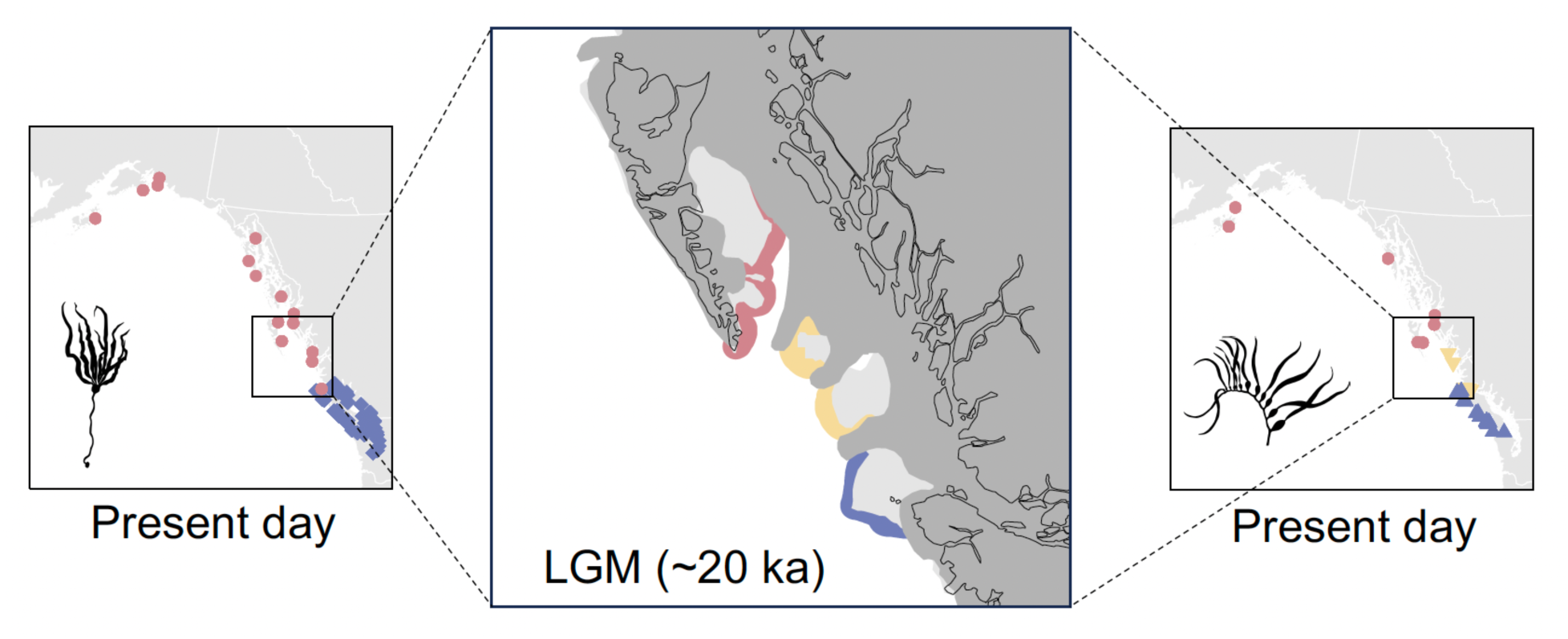
Hypothesized refugial scenario during the Last Glacial Maximum (LGM), showing a qualitatively interpreted summary of all results, especially Fig. 6. The central panel shows hypothesized refugia off the coasts of southern Haida Gwaii (red), central British Columbia (yellow), and northern Vancouver Island (blue), with populations sampled in the present day that originated primarily from these refugia in corresponding colours for *Nereocystis* (left) and *Macrocystis* (right). The central panel depicts LGM glaciers (dark grey), exposed continental shelf (light grey), and the modern coastline (black lines). The location of glaciers and continental shelf is a cartoon representation redrawn by hand from Shaw *et al*. (2020).

### Kelp persistence in northern refugia

Descriptive phylogeographic approaches and the *Spacetrees* model agreed in identifying northern refugia along glacial margins, providing strong support for survival of both species in ice-free pockets of central and northern BC. Supporting evidence from descriptive approaches includes (1) higher genetic diversity in this region than elsewhere in AKBCWA (Fig. 3); (2) pairwise directionality index (*ψ*) values (Fig. 4) suggesting that range expansion proceeded from this region in both a northerly and southerly direction; and (3) simple ENMs suggesting that low- to moderate-quality habitat was available in this region (Supporting Information Figs. S6, S7).

However, although these lines of evidence are strongly compatible with northern refugia, qualitative interpretation of genetic diversity and *ψ* does not provide definitive evidence of refugial locations for several reasons. High genetic diversity is expected in refugial locations due to loss of genetic diversity from successive founder events during postglacial expansion (Hewitt, 2000), but patterns of genetic diversity can have multiple causes. High genetic diversity may occur in areas of secondary contact between lineages expanding from other areas (Petit *et al*., 2003). Alternatively, genetic diversity may reflect contemporary population size, which varies widely and is highly correlated with *π* in both species (Bemmels *et al*., 2025). In addition, the core-periphery hypothesis predicts that genetic diversity should be higher in the middle of a species’ range and lower at the edges due to decreased gene flow and stronger genetic drift in small populations at the range margins (Eckert *et al*., 2008). Core-periphery dynamics may also severely bias *ψ* (Kemppainen *et al*., 2024). As AKBCWA could be considered a contiguous biogeographic unit given low abundance farther south in Oregon (Supporting Information Fig. S4), the north coast of BC would represent the geographic core of this unit and correspond to the area of expected highest diversity. However, the core-periphery hypothesis has limited support in other marine taxa (Cárcamo *et al*., 2025) and would require that the genetic signal of postglacial range expansion has been totally erased, which seems unlikely given the short time that has elapsed since the LGM and strong genetic structure (Fig. 1 and Supporting Information Fig. S2). Finally, as expected immediately following range expansion (Kemppainen *et al*., 2024),*ψ* was highly correlated with pairwise Δ*π* (inset plots in Fig. 4). Thus, *ψ* cannot be taken as an independent source of information to strengthen confidence in interpretations of *π,* but instead provides a visualization of the shared signal inherent in both statistics.

Given the difficulties in qualitative interpretation of genetic patterns, the *Spacetrees* analysis that leveraged spatial signatures from >800 local trees across the genome to infer the coordinates of genetic ancestors provided a powerful tool to locate refugia and to visualize key features of postglacial range expansion. *Spacetrees* showed increasing geographic convergence of ancestors backward in time toward north-central BC in both species (Fig. 6). At 20 ka, there is a broad spatial distribution of inferred ancestor locations (90% contour lines; Fig. 6). However, as coalescence predates the time of population splitting in the absence of gene flow (Edwards & Beerli, 2000), some lineages that expanded from the same refugium may have coalesced far earlier than 20 ka. It is thus relevant to consider trajectories of ancestor locations prior to 20 ka as coalescence continues. At 40 ka in *Nereocystis* and 40-80 ka in *Macrocystis*, mean ancestral locations were mostly distributed from northern Vancouver Island to Haida Gwaii (Fig. 6), highlighting the strong signal of range expansion from this region. Importantly, there was no evidence of convergence toward the south in either species.

### A region of multiple refugia

In addition to confirming the likely existence of northern refugia in north-central BC, ARG-based analyses point to the potential existence of multiple refugia within this region (summarized in Fig. 7). Evidence of multiple refugia is clearest in *Macrocystis*, where the mean ancestor locations for the seven genetic clusters converge toward three separate geographic locations (Fig. 6b): southern Haida Gwaii, the central coast of BC, and northern Vancouver Island. This convergence to three locations is not transitory, as might be expected if it reflected an intermediate state of migration between a refugial and non-refugial scenario, but clearly persists from 20-80 ka (Fig. 6b).

Furthermore, population divergence times in *Macrocystis* are more recent than the LGM between populations that converge toward the same location, but mostly overlap with or are more ancient than the LGM between populations that converge toward different locations (Fig. 5d and Supporting Information Table S5), suggesting that there was more than one refugium present at the LGM. Additionally, bottlenecks with minimal *N*_e_ in the thousand to tens of thousands (Fig. 5b) suggest at least moderately sized refugia and are not compatible with collapse of all populations into a single, extremely small population. Finally, the division of populations into three groups along the first PC axis (Fig. 1f) corresponds precisely to the division of populations into three ancestral locations in *Spacetrees* (Fig. 6b), suggesting that contemporary genetic structure may have its origins in the three distinct LGM refugia.

In *Nereocystis*, the number of refugia is more difficult to determine. Ancestor locations show less spatial convergence and lower stability over time, but spatial convergence from 20-40 ka (Fig. 6a) suggests at least two main refugia in southern Haida Gwaii and northern Vancouver Island (summarized in Fig. 7). In addition, divergence times between northern and southern populations are more ancient than the LGM (Fig. 5c and Supporting Information Table S5), as expected if northern and southern populations are derived from separate refugia. The ancestors of the northernmost (light blue) cluster are inferred to be located at latitudes equivalent to Southeast Alaska from 20-40 ka (Fig. 6a), but it is unlikely there was a separate refugium in Alaska because the inferred divergence time between the two northernmost clusters dates to the LGM (Fig. 5c) and the 90% contour line for this cluster at 20 ka includes southern areas, indicating some coalescence with southern populations. Instead, we speculate that this genetic cluster may have originated through long-distance migration across the Gulf of Alaska, which the Brownian motion dispersal model in *Spacetrees* is not designed to accommodate (Osmond & Coop, 2024).

Remarkably, the inferred refugial locations correspond almost exactly with geological reconstructions suggesting several disjunct ice-free areas of coastline separated by glacial lobes (Shaw *et al*., 2020; Mann & Gaglioti, 2024). These ice-free areas (Fig. 7) were located off the coast of southeastern Haida Gwaii (the “Hecate refugium”), the central coast of BC (though there were likely two major ice-free pockets along the central coast whereas our genetic analyses inferred only one), and northern Vancouver Island, precisely where *Spacetrees* inferred refugia for *Macrocystis* and most populations of *Nereocystis*. As the spatial spread of inferred ancestral locations was extremely broad at all time periods (Fig. 6), suggesting high uncertainty, we cannot rule out that this result could have been coincidence. Nonetheless, ENM projections largely suggest that LGM persistence in these ice-free regions is plausible (Supporting Information Figs. S6,S7). Although suitable habitat was not predicted in this region for *Macrocystis* in the ensemble climate model excluding CCSM3 (Supporting Information Fig. S6b), the ensemble model does not correctly account for increased ocean salinity due to freshwater glacier formation and a drop in sea level during the LGM (Sbrocco, 2014). As salinity was an important predictor of habitat suitability for *Macrocystis* (Supporting Information Table S3), the CCSM3 model that corrects for increased salinity may be most appropriate for this species. Additional areas of predicted suitability exist in southern Vancouver Island and Southeast Alaska but were unlikely to have contained refugia for our study species. Southern Vancouver Island experienced an extremely late local LGM (Mann & Hamilton, 1995; Mann & Gaglioti, 2024) and was eventually overrun by glaciers, which is not reflected in the glacial reconstruction for the global LGM (Gillespie *et al*., 2004) depicted in Supporting Information Figs. S6 and S7. Meanwhile, seasonal sea ice may have covered the entire coast of Southeast Alaska during the LGM (Praetorius *et al*., 2023; Mann & Gaglioti, 2024) suggesting that this region presented very different winter environmental conditions than either species experiences today, which casts doubt on inferred habitat suitability north of Haida Gwaii.

### Global relationships in Macrocystis

*Macrocystis* populations from different global regions have a long and independent history. Divergence times from *Relate* (Fig. 5c) suggest that the earliest split occurred between the Northern and Southern Hemispheres at 2.3 Ma, near the onset of the Pleistocene (2.58 Ma). The late Pliocene to early Pleistocene was a period of major transition, with Pleistocene oceans characterized by colder temperatures and strengthened cold-water upwelling in the Pacific (Filippelli & Flores, 2009). Closure of the Isthmus of Panama ∼2.75 Ma also reorganized ocean circulation and resulted in increased nutrient concentrations at low latitudes of the eastern Pacific (Schneider & Schmittner, 2006). We speculate that these global changes could have created the conditions necessary for dispersal to the South Pacific from the North Pacific, where *Macrocystis* likely originated (Coyer *et al*., 2001; Starko *et al*., 2019; Assis *et al*., 2023). Dispersal may have been most likely during a glacial period when distributions of temperate seaweed species shifted toward lower latitudes (Song *et al*., 2021), and perhaps could have made use of (sub)tropical islands as stepping stones (Assis *et al*., 2018). Deep divergences also exist within the Southern Hemisphere (619 ka split between Northern and Southern Chile), in agreement with previous findings (Lindstrom, 2023; Gonzalez *et al*., 2023).

Within North America, divergences between morphs and between California and AKBCWA are more recent but substantially predate the LGM (Fig. 5c). However, several lines of evidence hint that there may have been some genetic contact between California and AKBCWA after they initially diverged, including (1) the intermediate phylogenetic (Supporting Information Fig. S3a) and PCA (Fig. 2) position of California *integrifolia* between AKBCWA and California *pyrifera*; (2) reduced *d*_XY_ between California *integrifolia* and southern AKBCWA, but not northern AKBCWA (Supporting Information Fig. S2); and (3) signal of gene flow between California *integrifolia* and southern BC populations (Box 2 in Supporting Information Fig. S3c).

Interpreting the phylogenetic placement of California *integrifolia* and divergence times with other populations at face value would suggest that California *integrifolia* and AKBCWA populations share a single origin, and there has been subsequent gene flow between California *integrifolia* and southern AKBCWA (Hypothesis 1; Supporting Information Fig. S3b). Alternatively, it is also plausible that California *integrifolia* is sister to southern AKBCWA but has subsequently introgressed with northern California *pyrifera* (Hypothesis 2; Supporting Information Fig. S3b), causing an intermediate phylogenetic position (Fig. S3a). This scenario may have occurred, for example, if California *integrifolia* is descended from long-distant migrants from southern AKBCWA. We are unable to distinguish these two scenarios and the relationship between California and AKBCWA requires further study, including expanded sampling of *integrifolia* from multiple locations in California. Relationships among California *pyrifera* also require clarification, given the unexpected clustering of a northern Californian population from Bodega Bay (purple) with the distant Channel Islands (pink) in the PCA (Fig. 2), possibly suggesting long-distance migration and complex but undocumented genetic structure within California (Alberto *et al*., 2010; Johansson *et al*., 2015; Gonzalez *et al*., 2023; Assis *et al*., 2023). Furthermore, the close clustering of *pyrifera* morphs from Alaska with presumed *integrifolia* morphs from southern BC and Washington rather than with other *pyrifera* morphs from California (PC1 in Fig. 2b; phylogeny in Supporting Information, Fig. S3a) highlights how examining multiple geographic transition zones between morphs (i.e., both California and Alaska) will be required to fully understand the genetic basis of morph identity in North America (Gonzalez *et al*., 2023; Gonzalez & Raimondi, 2024) or possible lack thereof.

### Coastal ecosystems and human migration

The inferred LGM persistence of canopy-forming kelps along the central to northern coast of BC has implications that extend to other species, including humans. Because these kelps are foundation species that create upright, spatially-structured habitats upon which numerous species rely (Teagle *et al*., 2017; Wernberg *et al*., 2019), their presence during the LGM suggests that kelp forest ecosystems could plausibly have persisted at these northern latitudes throughout the Last Glacial Period. Although phylogeographic histories for most coastal marine species from this region are poorly understood (Marko *et al*., 2010), numerous other species of algae (Lindstrom *et al*., 1997, 2021, 2025), fish (Hickerson & Ross, 2001; Smith *et al*., 2001; Hickerson & Cunningham, 2005; Marko *et al*., 2010), and invertebrates (Marko *et al*., 2010) may also have survived glaciation in northern refugia along the BC and Southeast Alaska coast, hinting that this region may have hosted biodiverse kelp forests and other coastal marine ecosystems during the LGM. However, further study of diverse organisms will be needed to confirm this hypothesis, as co-distributed species often exhibit species-specific phylogeographic histories (Stewart *et al*., 2010; Papadopoulou & Knowles, 2015) and communities with no close modern analogues were common in the past (Jackson & Williams, 2004; Graham, 2005). In Southern California, kelp forest productivity is believed to have fluctuated since the LGM, peaking during the mid-Holocene (Graham *et al*., 2010). It is thus possible that the LGM kelp forests of north-central BC may have substantially differed from modern kelp forests in their species assemblages or ecosystem functions.

Nonetheless, kelp forest presence in north-central BC during the LGM provides a key piece of evidence in support of the Kelp Highway Hypothesis (Erlandson *et al*., 2007) of the peopling of the Americas. It was once widely believed that humans first migrated to the Americas over the Bering Land Bridge and through an ice-free corridor that opened in the middle of the continent as glaciers melted (Meltzer, 1993). This model has now been largely rejected in favour of the Kelp Highway Hypothesis (Braje *et al*., 2017), which suggests migration along the coast of Alaska and BC by people relying on the abundant resources provided by kelp forests (Erlandson *et al*., 2007) and other coastal marine and terrestrial ecosystems (Erlandson *et al*., 2015). This migration would have occurred no earlier than ∼17 ka, once deglaciation opened a continuous route through Southeast Alaska (Erlandson *et al*., 2007; Lesnek *et al*., 2018; Praetorius *et al*., 2023). Definitive human archaeological evidence of the Kelp Highway Hypothesis is lacking (Braje *et al*., 2017), but it depends critically on the existence of a glacier-free coastline with appropriate ecological conditions to support human populations (Lesnek *et al*., 2018). Although migrating populations may have used additional coastal resources not associated with kelp forests (Erlandson *et al*., 2015), perhaps even hunting offshore from marine sea ice during colder months of the year (Lesnek *et al*., 2018; Praetorius *et al*., 2023), the familiarity of early Americans with kelp forests is strongly suggested by their early, rapid expansion along the Pacific Rim (Braje *et al*., 2017) to areas that today support *Macrocystis* forests (Macaya & Zuccarello, 2010), and evidence of human use of *Macrocystis* in southern Chile as early as ∼14 ka (Dillehay *et al*., 2008). However, arguments to date for the importance of kelp forests along the putative coastal migration route have largely been inspired by comparisons with modern ecosystems (Erlandson *et al*., 2007, 2015) rather than reconstructions of the late-Pleistocene distribution of kelp. In contrast, our inference of refugia along the formerly glaciated coastlines of northern and central BC suggests canopy-forming kelp forests may have already existed during the LGM at relatively high latitudes from which they could have rapidly expanded as glaciers melted, thus helping to facilitate the arrival of North America’s earliest peoples.

## Acknowledgments

Alaska: Collections of *Macrocystis* were facilitated by Susan Saupe (Cook Inlet Regional Citizens Advisory Council). | Canada: We thank the Gitga’at, Gitxaała, Haida, Haisla, Heiltsuk, Kitasoo-Xai’xais, Kitselas, Kitsumkalum, K’ómoks, Mamalilikulla, Metlakatla, Tlowitsis, and Wei Wai Kum First Nations for permission to reanalyze DNA sequences from samples collected from their territories. For a BioCultural Notice of cultural rights and responsibilities regarding these samples see (Bemmels *et al*., 2025). Funding was provided by the Natural Sciences and Engineering Research Council of Canada (RGPIN-2021-02482 to GLO; RGPIN–2021-03207 to MMO) and Genome British Columbia (GIRAFF Grant to GLO). | California: This study is a contribution of the Marine Networks Consortium (PIs: Michael N. Dawson, Rachael A. Bay) as part of the California Conservation Genomics Project (PI: H. Bradley Shaffer), with funding provided to the University of California by the State of California, State Budget Act of 2019 (UC Award ID RSI-19-690224). We thank Brenda Cameron for help with extraction and preparation of genomic libraries. We also thank the following individuals for help with collection: Shannon Myers, Jason Toy, Emily Donham, Zoe Scholtz, Chelsea Williams, Vaness Guerra, Alexis Necarsulmer, Jon Powl Ewing, Adam and Jenesa Wall, and Angela Korabik.

## Author contributions

Conceptualization, JBB, GLO; formal analysis, JBB, MMO; funding acquisition, RAB, SCL, MMO, GLO; investigation, JBB; resources, KJK, SRP, RAB, KMG, SCL; software, MMO; supervision, GLO; writing – original draft, JBB; writing – review and editing, GLO with contributions from all authors.

## Data availability

Raw DNA sequences will be made openly available in the NCBI SRA at https://www.ncbi.nlm.nih.gov/sra upon manuscript acceptance, BioProject Numbers PRJNA1256528, PRJNA1140647, PRJNA1164249, PRJNA938791, PRJNA661280, and PRJEB55054. Custom scripts used to process data will be made openly available at Zenodo upon manuscript acceptance.

## Supporting information

**Fig. S1** Map of geographic features mentioned in the text

**Fig. S2** Genetic divergence and differentiation

**Fig. S3** Phylogenetic relationships and gene flow among global *Macrocystis*

**Fig. S4** Occurrence records used in ecological niche models

**Fig. S5** Habitat suitability in the current time period

**Fig. S6** Habitat suitability during the Last Glacial Maximum – ensemble model

**Fig. S7** Habitat suitability during the Last Glacial Maximum – CCSM3 model

**Table S1** Sampling information for newly sequenced data

**Table S2** Sampling information for previously published sequencing data

**Table S3** Ecological niche model predictor variable importance metrics

**Table S4** Ecological niche model performance metrics

**Table S5** Divergence time estimates between populations

**Methods S1** Methodological details

