## Supporting Information for "Spatial inference of ancestor locations suggests northern refugia for canopy-forming kelps in the Northeast Pacific"

**Fig. S1** Map of geographic features mentioned in the text

**Fig. S2** Genetic divergence and differentiation

**Fig. S3** Phylogenetic relationships and gene flow among global *Macrocystis*

**Fig. S4** Occurrence records used in ecological niche models

**Fig. S5** Habitat suitability in the current time period

**Fig. S6** Habitat suitability during the Last Glacial Maximum – ensemble model

**Fig. S7** Habitat suitability during the Last Glacial Maximum – CCSM3 model

**Table S1** Sampling information for newly sequenced data

**Table S2** Sampling information for previously published sequencing data

**Table S3** Ecological niche model predictor variable importance metrics

**Table S4** Ecological niche model performance metrics

**Table S5** Divergence time estimates between populations

**Methods S1** Methodological details

**Supporting references**

**Fig. S1** Map of geographic features mentioned in the text, showing the Northeast Pacific coastline of Alaska, British Columbia, and Washington. Terrestrial features are labelled in black text and marine features in blue. General regions of the coastline that we refer to but which do not have formal names or precisely defined boundaries are labelled in green text.

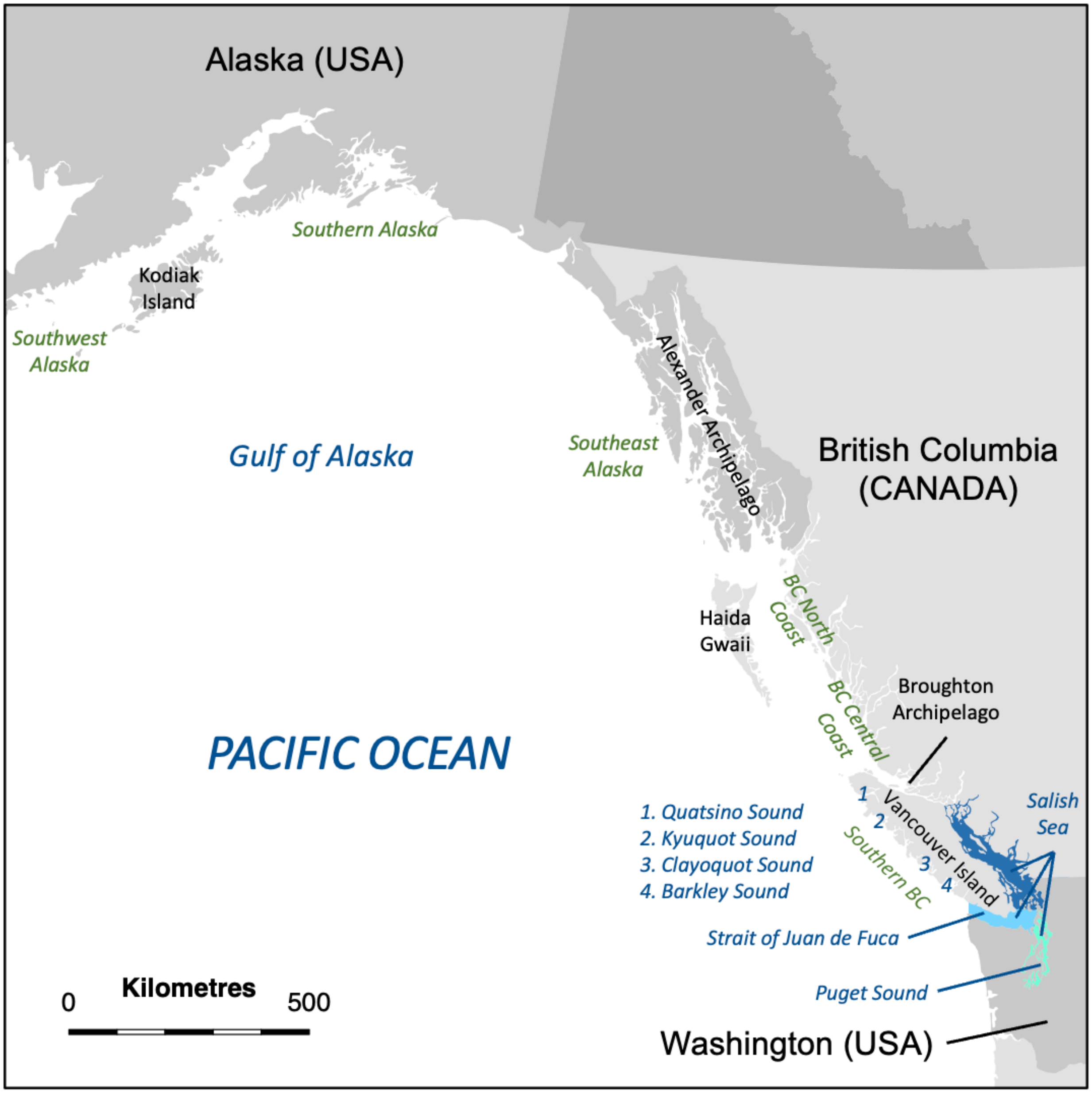

**Fig. S2** Genetic divergence and differentiation among genetic clusters and geographic regions of (a) *Nereocystis* and (b) *Macrocystis*. Above the diagonal: genetic divergence ($d_{XY}$); along the diagonal: nucleotide diversity ($\pi$); below the diagonal: genetic differentiation ($F_{ST}$). Genetic clusters are indicated by labels and colours corresponding to Fig. 1, with additional global regions not included in genetic clustering analyses in grey. White shading indicates that $F_{ST}$ could not be reliably calculated as only a single individual was available for one or more of the geographic regions in the comparison. AK: Alaska; arch: archipelago; CA: California.

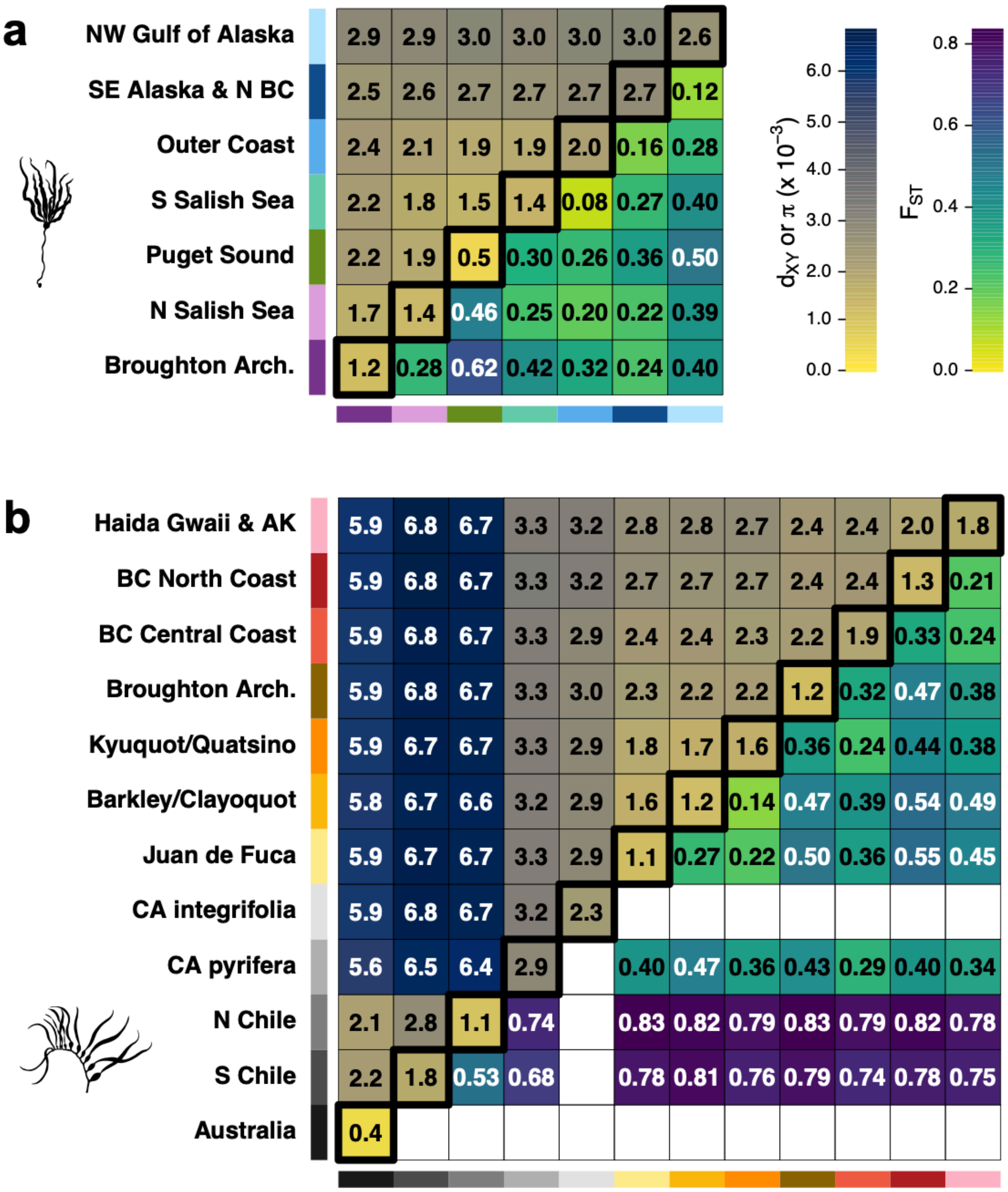

**Fig. S3** Phylogenetic relationships and gene flow among global *Macrocystis* populations. (a) Phylogeny constructed in *IQ-TREE*. Branches represent individuals, with one individual selected per population. Vertical bars identify the genetic clusters (colours) or global regions (greyscale) to which individuals belong. Branch lengths are scaled in units of nucleotide substitutions per site. AK: Alaska; arch: archipelago; CA: California; SH-aLRT: SH-like approximate likelihood ratio test. (b) Competing hypotheses about the phylogenetic position of the *integrifolia* morph from northern California in relation to other regions of North America. Red arrows represent gene flow. (c) Tests of gene flow among North American populations inferred in *Dsuite.* Branches of the phylogenies represent populations (and dotted blue lines represent common ancestors), with bars at branch tips coloured according to genetic clusters and regions from (a). Shading of cells corresponds to the magnitude of the *f*-branch ($f_b$) statistic, representing gene flow among branches of the phylogeny. White cells indicate that $f_b$ was not statistically significant and inferences were not possible in grey cells. The *integrifolia* morph from northern California is underlined in red, with highlighted black Box 1 indicating gene flow between *integrifolia* and northern California *pyrifera*, and Box 2 between *integrifolia* and southern AKBCWA populations.

[[Figure displayed on the next page.]]

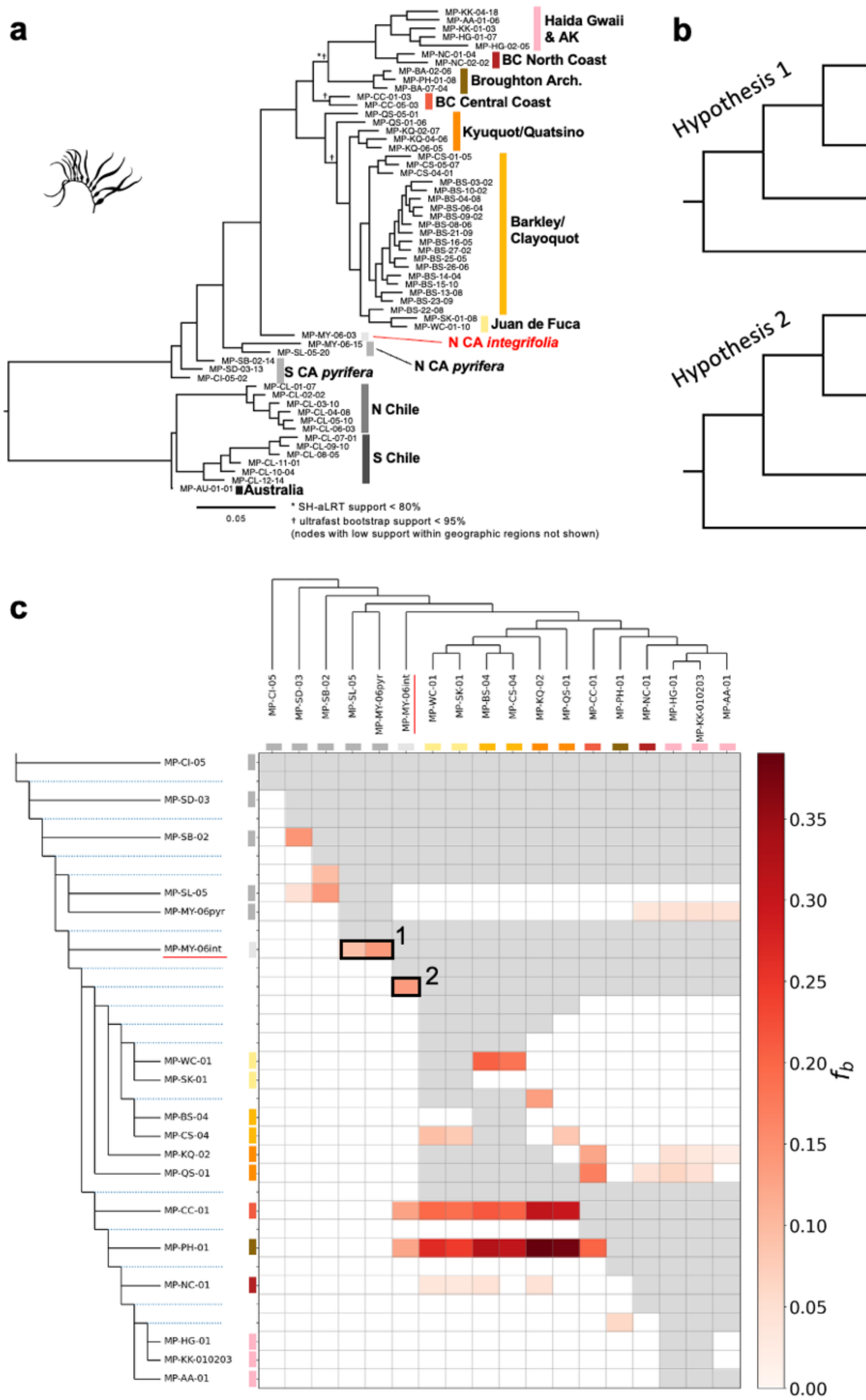
a
Haida Gwaii & AK
BC North Coast
Broughton Arch.
BC Central Coast
Kyuquot/Quatsino
Barkley/ Clayoquot
Juan de Fuca
N CA integrifolia
N CA pyrifera
S CA pyrifera
N Chile
S Chile
Australia
0.05
* SH-aLRT support < 80%
† ultrafast bootstrap support < 95%
(nodes with low support within geographic regions not shown)
b
Hypothesis 1
North AKBCWA
South AKBCWA
North CA integrifolia
North CA pyrifera
South CA pyrifera
Hypothesis 2
North AKBCWA
South AKBCWA
North CA integrifolia
North CA pyrifera
South CA pyrifera
c
MP-CI-05
MP-SD-03
MP-SB-02
MP-SL-05
MP-MY-06pyr
MP-MY-06int
MP-WC-01
MP-SK-01
MP-BS-04
MP-CS-04
MP-KQ-02
MP-QS-01
MP-CC-01
MP-PH-01
MP-NC-01
MP-HG-01
MP-KK-010203
MP-AA-01
1
2
$f_b$
0.35
0.30
0.25
0.20
0.15
0.10
0.05
0.00

**Fig. S4** Filtered occurrence records used in ecological niche models (ENMs). (a) *Nereocystis* occurrences. (b) *Macrocystis* occurrences. (c) Brown algae occurrences used as background points.

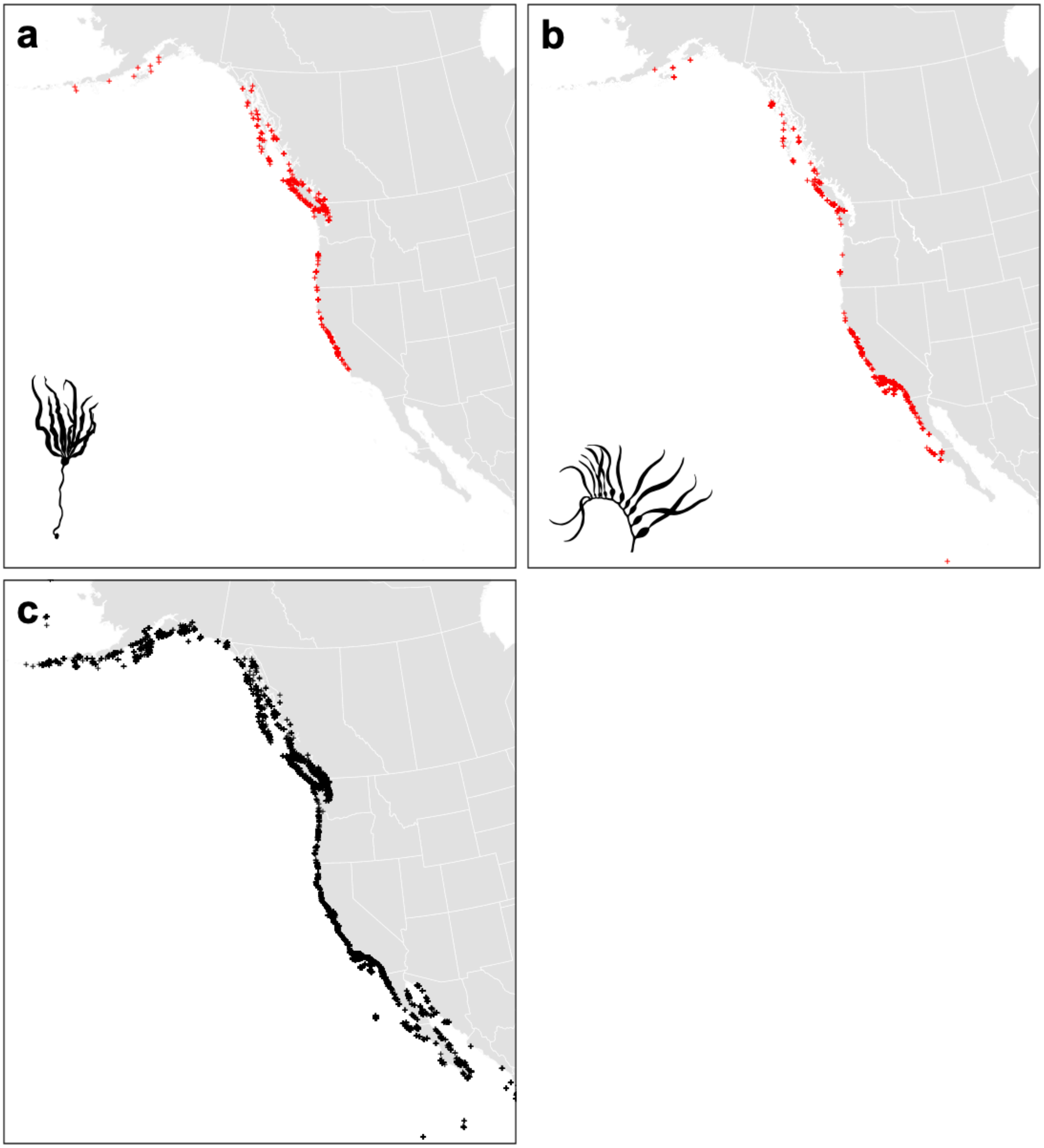

**Fig. S5** Predicted habitat suitability from ENMs in the current time period for (a) *Nereocystis* and (b) *Macrocystis*. White pixels represent areas predicted to be unsuitable according to species-specific thresholds that retain 95% of unique true occurrences.

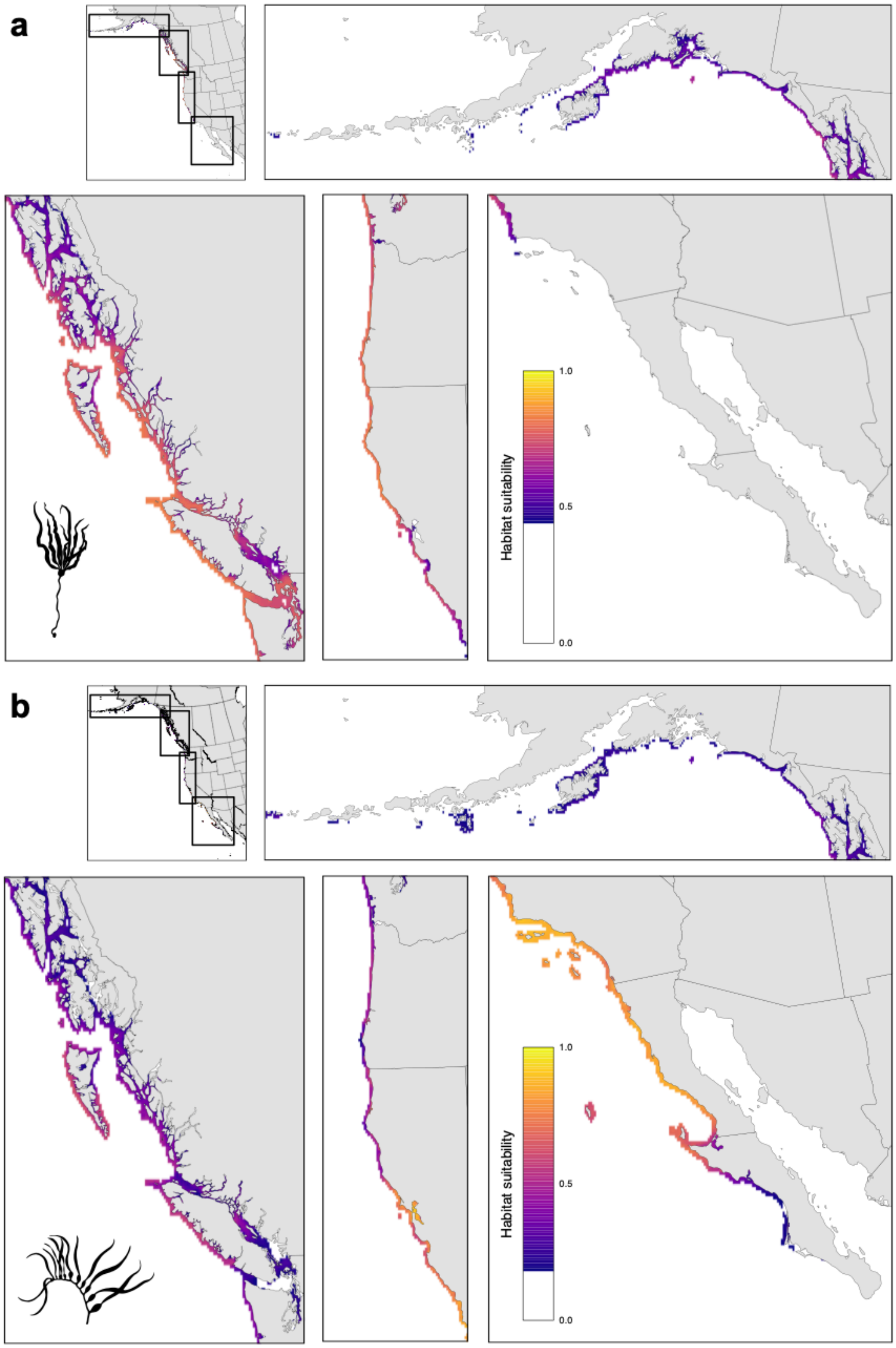

**Fig. S6** Predicted habitat suitability from ENMs during the Last Glacial Maximum (LGM) according to the ensemble of all climate models except CCSM3, for (a) *Nereocystis* and (b) *Macrocystis*. White pixels represent areas predicted to be unsuitable according to species-specific thresholds that retain 95% of unique true occurrences. Dark grey areas represent the maximum extent of glaciation (Gillespie *et al.*, 2004).

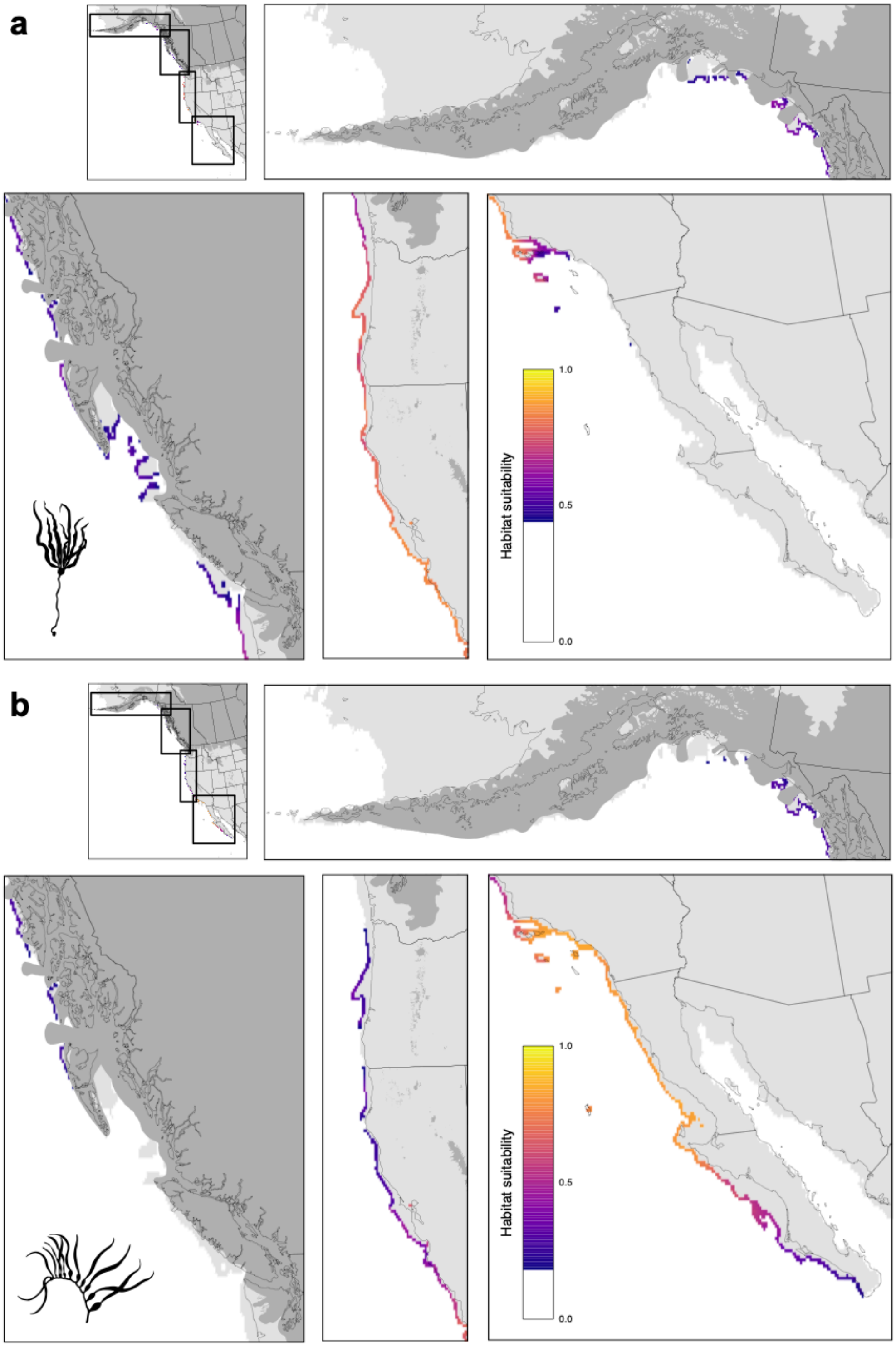

**Fig. S7** Predicted habitat suitability from ENMs during the Last Glacial Maximum (LGM) according to CCSM3 climate model, for (a) *Nereocystis* and (b) *Macrocystis*. White pixels represent areas predicted to be unsuitable according to species-specific thresholds that retain 95% of unique true occurrences. Dark grey areas represent the maximum extent of glaciation (Gillespie *et al.*, 2004).

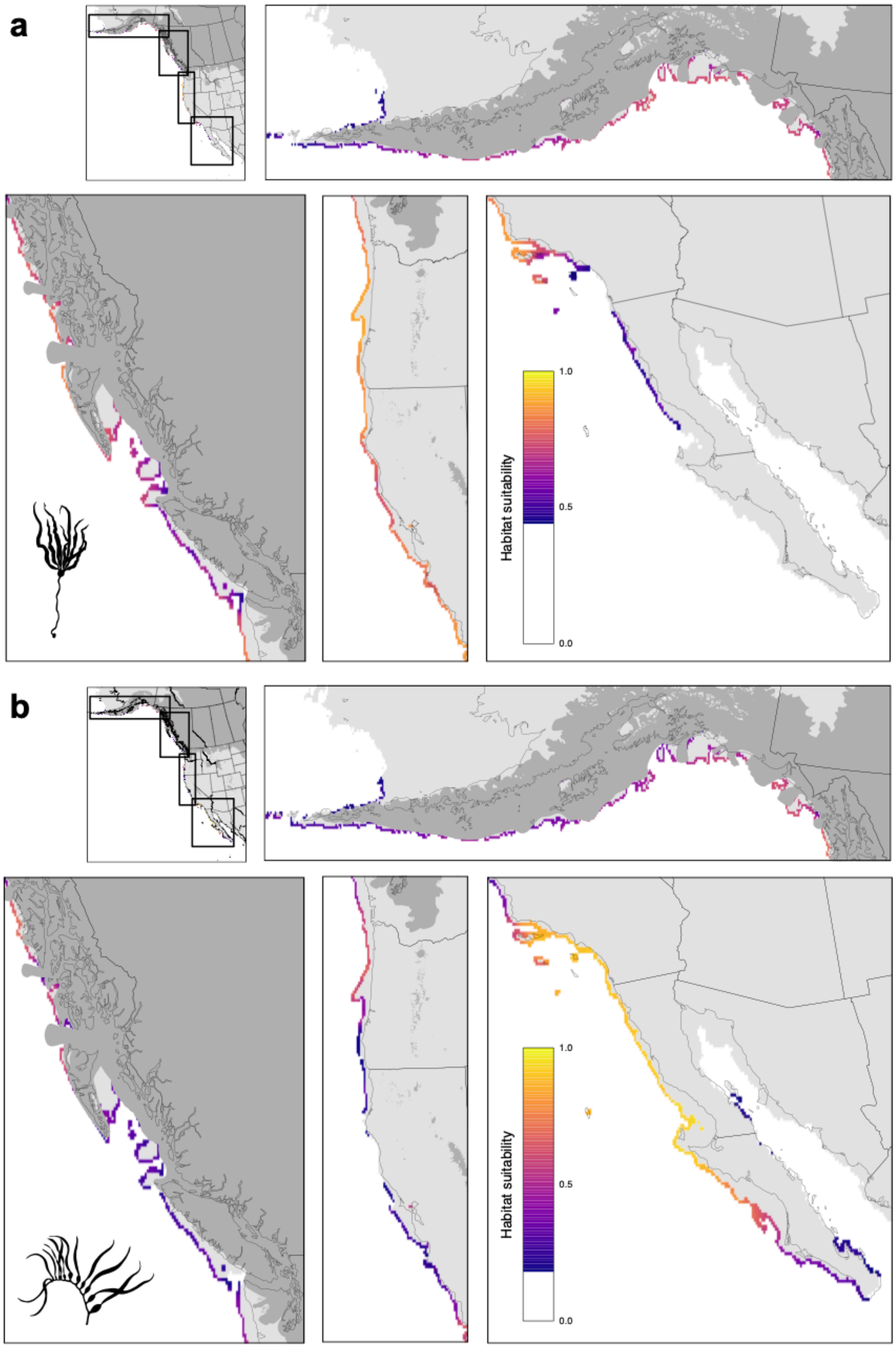

**Table S1** Collecting localities and sample sizes for newly sequenced populations. Sp.: species; N: Nereocystis; M: Macrocystis; Jur.: political jurisdiction; AK: Alaska; CA: California; pyr, pyrifera morph; unk, morph identity unassessed; nseq: number of individuals sequenced; n1x: number of individuals retained in the 1x dataset (see Materials and methods) after filtering; n8x: number of individuals retained in the 8x dataset after filtering; accession: NCBI BioProject Accession Number.

| Sp. | Jur. | Site code | Site name | Morph | Latitude | Longitude | $n_{seq}$ | $n_{1x}$ | $n_{8x}$ | Accession |
|---|---|---|---|---|---|---|---|---|---|---|
| N | AK | NL-AA-02 | Vallenar Rock | n/a | 55.38353 | -131.87645 | 7 | n/a | 4 | PRJNA1256528 |
| N | AK | NL-AA-03 | Hayward Strait | n/a | 57.14853 | -135.55962 | 7 | n/a | 3 | PRJNA1256528 |
| N | AK | NL-AA-04 | Lynn Sisters | n/a | 58.42288 | -135.087 | 7 | n/a | 6 | PRJNA1256528 |
| N | AK | NL-AA-05 | Little Port Walter | n/a | 56.385 | -134.641 | 7 | n/a | 2 | PRJNA1256528 |
| N | AK | NL-KK-05 | Woody Island | n/a | 57.7647 | -152.3514 | 7 | n/a | 6 | PRJNA1256528 |
| N | AK | NL-PW-01 | Shelter Bay | n/a | 60.4258 | -146.6828 | 7 | n/a | 7 | PRJNA1256528 |
| N | AK | NL-PW-02 | Tatitlek Narrows | n/a | 60.84743 | -146.69668 | 7 | n/a | 6 | PRJNA1256528 |
| N | AK | NL-PW-03 | Fox Farm Bay | n/a | 59.9664 | -148.1513 | 7 | n/a | 6 | PRJNA1256528 |
| M | AK | MP-AA-01 | Salisbury Sound Site 1 | pyr | 57.35 | -135.77 | 7 | 7 | 7 | PRJNA1256528 |
| M | AK | MP-KK-01 | NW Afognak Island Site 1 | pyr | 58.405 | -152.82917 | 3 | 3 | 3 | PRJNA1256528 |
| M | AK | MP-KK-02 | NW Afognak Island Site 2 | pyr | 58.40472 | -152.82694 | 2 | 2 | 2 | PRJNA1256528 |
| M | AK | MP-KK-03 | NW Afognak Island Site 3 | pyr | 58.40772 | -152.79644 | 2 | 2 | 2 | PRJNA1256528 |
| M | AK | MP-KK-04 | Kiliuda Bay | pyr | 57.3 | -152.91 | 7 | 7 | 7 | PRJNA1256528 |
| M | CA | MP-CI-01 | Blue Cavern | unk | 33.44844 | -118.47891 | 5 | 3 | 0 | PRJNA1140647 |
| M | CA | MP-CI-02 | Ironbound Cove | unk | 33.44768 | -118.57765* | 10 | 10 | 0 | PRJNA1140647 |
| M | CA | MP-CI-03 | Two Harbors | unk | 33.44549 | -118.49831 | 1 | 1 | 0 | PRJNA1140647 |
| M | CA | MP-CI-04 | Two Harbors, beach | unk | 33.42659 | -118.50619 | 5 | 5 | 0 | PRJNA1140647 |
| M | CA | MP-LA-01 | Malibu | unk | 34.02193 | -118.77394 | 6 | 6 | 0 | PRJNA1140647 |
| M | CA | MP-LA-02 | KOU Rock | unk | 33.7203 | -118.33722 | 6 | 6 | 0 | PRJNA1140647 |
| M | CA | MP-LA-03 | Palos Verdes, lower limit | unk | 33.70694 | -118.28589 | 6 | 6 | 0 | PRJNA1140647 |
| M | CA | MP-LA-04 | Recreation Point | unk | 33.54213 | -117.795 | 6 | 6 | 0 | PRJNA1140647 |
| M | CA | MP-LA-05 | Laguna Beach | unk | 33.52987 | -117.78093 | 5 | 5 | 0 | PRJNA1140647 |
| M | CA | MP-MY-01 | Terrace Point | unk | 36.94487 | -122.06428 | 9 | 9 | 0 | PRJNA1140647 |
| M | CA | MP-MY-02 | Stillwater Cove | unk | 36.56012 | -121.94732 | 5 | 5 | 0 | PRJNA1140647 |
| M | CA | MP-MY-03 | Cooper Point | unk | 36.26315 | -121.85662 | 6 | 6 | 0 | PRJNA1140647 |
| M | CA | MP-MY-04 | Landels-Hill Big Creek Reserve | unk | 36.06888 | -121.601 | 5 | 5 | 0 | PRJNA1140647 |
| M | CA | MP-MY-05 | Mill Creek | unk | 35.97976 | -121.49046 | 6 | 6 | 0 | PRJNA1140647 |
| M | CA | MP-RC-01 | Bodega Bay | unk | 38.31320 | -123.0514 | 8 | 8 | 0 | PRJNA1140647 |
| M | CA | MP-SB-01 | Mohawk Reef/Mesa Lane Beach | unk | 34.39456 | -119.73 | 10 | 10 | 0 | PRJNA1140647 |
| M | CA | MP-SD-01 | Boomer Beach | unk | 32.8522 | -117.276 | 4 | 4 | 0 | PRJNA1140647 |
| M | CA | MP-SD-02 | La Jolla | unk | 32.80959 | -117.28662 | 10 | 10 | 0 | PRJNA1140647 |
| M | CA | MP-SL-01 | Hazards | unk | 35.28 | -120.88 | 6 | 6 | 0 | PRJNA1140647 |
| M | CA | MP-SL-02 | San Luis | unk | 35.24408 | -120.9011 | 6 | 6 | 0 | PRJNA1140647 |
| M | CA | MP-SL-03 | Point Buchon | unk | 35.24124 | -120.8955 | 6 | 6 | 0 | PRJNA1140647 |
| M | CA | MP-SL-04 | Shell Beach | unk | 35.16902 | -120.69715 | 6 | 6 | 0 | PRJNA1140647 |

* The original longitude recorded as -118.51165 for Ironbound Cove occurs on land and was assumed to be a transcription error. The site was georeferenced to Iron Bound Bay at longitude -118.57765.

**Table S2** Collecting localities and sample sizes for populations from previously published data. Sp.: species; N: Nereocystis; M: Macrocystis; Jur.: political jurisdiction; BC: British Columbia; WA: Washington; CA: California; CL: Chile; AU: Australia; unk, morph identity unassessed; int, integrifolia morph; pyr, pyrifera morph; nseq: number of individuals sequenced; n1x: number of individuals retained in the 1x dataset (see Materials and methods) after filtering; n8x: number of individuals retained in the 8x dataset after filtering; accession: NCBI BioProject Accession Number; B, Bemmels et al. (2025); G, Gonzalez et al. (2023); M, Molano et al. (2022); I, Iha et al. (2023).

| Sp. | Jur. | Site code | Site name | Morph | Latitude | Longitude | $n_{seq}$ | $n_{1x}$ | $n_{8x}$ | Accession | Citation |
|---|---|---|---|---|---|---|---|---|---|---|---|
| N | BC | various* | see Bemmels et al. (2025) | n/a | various | various | 330 | n/a | 314 | PRJNA1164249* | B |
| N | WA | various | see Bemmels et al. (2025) | n/a | various | various | 119 | n/a | 116 | PRJNA1164249 | B |
| M | BC | various* | see Bemmels et al. (2025) | unk,int† | various | various | 234 | 208 | 207 | PRJNA1164249* | B |
| M | WA | various | see Bemmels et al. (2025) | int† | various | various | 5 | 5 | 5 | PRJNA1164249 | B |
| M | CA | MP-MY-06int | Stillwater Cove (*integrifolia*) | int | 36.56506 | -121.94355 | 5 | 1 | 1 | PRJNA938791 | G |
| M | CA | MP-MY-06pyr | Stillwater Cove (*pyrifera*) | pyr | 36.56506 | -121.94355 | 5 | 5 | 5 | PRJNA938791 | G |
| M | CA | MP-SL-05 | Cayucos | pyr | 35.44646 | -120.93239 | 5 | 5 | 5 | PRJNA938791 | G |
| M | CA | MP-SB-02 | Arroyo Quemado | pyr | 34.47 | -120.119 | 16 | 12 | 1 | PRJNA661280 | M |
| M | CA | MP-CI-05 | Catalina Island | pyr | 33.462 | -118.511 | 16 | 16 | 1 | PRJNA661280 | M |
| M | CA | MP-SD-03 | Camp Pendleton | pyr | 33.157 | -117.36 | 17 | 17 | 6 | PRJNA661280 | M |
| M | CL | various | see Bemmels et al. (2025) | int,pyr | various | various | 60 | 58 | 58 | PRJNA938791 | G |
| M | AU | MP-AU-01 | Blackmans Bay | unk | -43.01733 | 147.32939 | 1 | 1 | 1 | PRJEB55054 | I |

* DNA sequences and associated metadata for samples provided by BC First Nations (Gitga'at, Gitxaała, Haida, Haisla, Heiltsuk, Kitasoo-Xai'xais, Kitselas, Kitsumkalum, K'ómoks, Mamalilikulla, Metlakatla, Tlowitsis, and Wei Wai Kum) are subject to a Biocultural Notice indicating cultural rights and responsibilities that require attention prior to data reuse. See Bemmels *et al.* (2025) for the full Biocultural Notice and further details.

† Although morph identity was not assessed, *Macrocystis* samples from southern BC and Washington are presumably of the *integrifolia* morph given that other morphs are not known to occur in this region (Macaya and Zuccarello, 2010; Gonzalez et al., 2023). Unassessed samples from northern BC (e.g., Haida Gwaii; Saunders & McDevit, 2014) may be either *integrifolia* or *pyrifera*.

**Table S3** Ecological niche model predictor variable importance metrics, mean (± standard deviation) across 20 resampling replicates. Each replicate contains 80% of the unique occurrences and background points. AUC (area under the receiver operating characteristic curve) metrics are for Jackknife tests of model performance with the focal variable excluded and with only the focal variable included, respectively. Permutation importance is the percentage decrease in training AUC when the values of the predictor variable are randomly permuted. SSS, sea surface salinity; SST sea surface temperature.

| Species | Predictor variable | AUC training without variable | AUC training with only variable | Permutation importance (%) |
|---|---|---:|---:|---:|
| *Nereocystis* | Mean annual SSS | 0.792 (±0.005) | 0.693 (±0.007) | 1.4 (±1.0) |
| | SST of the warmest month | 0.762 (±0.007) | 0.748 (±0.007) | 77.4 (±1.5) |
| | Annual range in SST | 0.750 (±0.008) | 0.566 (±0.009) | 21.2 (±1.6) |
| *Macrocystis* | Mean annual SSS | 0.795 (±0.006) | 0.722 (±0.007) | 22.0 (±2.5) |
| | SST of the warmest month | 0.755 (±0.006) | 0.751 (±0.007) | 56.7 (±2.1) |
| | Annual range in SST | 0.783 (±0.005) | 0.707 (±0.008) | 21.3 (±3.5) |

**Table S4** Ecological niche model performance metrics, mean (± standard deviation) across 20 resampling replicates. Each replicate contains 80% of the unique occurrences and background points. AUC, area under the receiver operating characteristic curve; TSS, true skill statistic.

| Species | AUC training | AUC testing | TSS training | TSS testing |
|---|---|---|---|---|
| *Nereocystis* | 0.797 (±0.007) | 0.781 (±0.009) | 0.536 (±0.013) | 0.541 (±0.009) |
| *Macrocystis* | 0.803 (±0.005) | 0.797 (±0.006) | 0.500 (±0.012) | 0.522 (±0.014) |

**Table S5** Relate divergence times (years) between populations from different genetic clusters (see also Fig. 5c,d). Median and range are calculated across all possible population pairs belonging to the two clusters. Sp.: species; N: Nereocystis; M: Macrocystis; B: between clusters; W: within clusters.

| Sp. | Type | Genetic cluster 1 | Genetic cluster 2 | Median divergence time | (Range) |
|---|---|---|---|---|---|
| N | B | NW Gulf of Alaska | Puget Sound | 119,475 | (85,984-119,475) |
| N | B | SE Alaska & N BC | Puget Sound | 85,984 | (44,535-119,475) |
| N | B | Broughton Arch. | NW Gulf of Alaska | 85,984 | (61,882-85,984) |
| N | B | Broughton Arch. | Puget Sound | 85,984 | (44,535-119,475) |
| N | B | S Salish Sea | NW Gulf of Alaska | 85,984 | (61,882-85,984) |
| N | B | SE Alaska & N BC | S Salish Sea | 61,882 | (11,948-85,984) |
| N | B | N Salish Sea | NW Gulf of Alaska | 61,882 | (61,882-85,984) |
| N | B | N Salish Sea | Puget Sound | 61,882 | (16,601-119,475) |
| N | B | Outer Coast | NW Gulf of Alaska | 61,882 | (32,052-85,984) |
| N | B | SE Alaska & N BC | N Salish Sea | 44,535 | (8,598-85,984) |
| N | B | SE Alaska & N BC | Broughton Arch. | 44,535 | (16,601-61,882) |
| N | B | Broughton Arch. | S Salish Sea | 44,535 | (32,052-61,882) |
| N | B | Broughton Arch. | Outer Coast | 44,535 | (32,052-61,882) |
| N | B | Outer Coast | Puget Sound | 38,293 | (11,948-85,984) |
| N | B | SE Alaska & N BC | Outer Coast | 32,052 | (50-61,882) |
| N | B | S Salish Sea | Puget Sound | 23,067 | (4,454-61,882) |
| N | B | SE Alaska & N BC | NW Gulf of Alaska | 16,601 | (8,598-44,535) |
| N | B | N Salish Sea | Broughton Arch. | 11,948 | (2,307-23,067) |
| N | B | N Salish Sea | Outer Coast | 11,948 | (4,454-32,052) |
| N | B | N Salish Sea | S Salish Sea | 8,598 | (4,454-23,067) |
| N | B | S Salish Sea | Outer Coast | 6,188 | (0-23,067) |
| N | W | SE Alaska & N BC | SE Alaska & N BC | 8,598 | (0-32,052) |
| N | W | S Salish Sea | S Salish Sea | 8,598 | (0-16,601) |
| N | W | NW Gulf of Alaska | NW Gulf of Alaska | 7,393 | (0-8,598) |
| N | W | Broughton Arch. | Broughton Arch. | 4,454 | (3,205-6,188) |
| N | W | Puget Sound | Puget Sound | 4,454 | (4,454-8,598) |
| N | W | N Salish Sea | N Salish Sea | 3,205 | (0-11,948) |
| N | W | Outer Coast | Outer Coast | 3,205 | (0-16,601) |
| M | B | Haida Gwaii & AK | Barkley/Clayoquot | 61,882 | (44,535-85,984) |
| M | B | Haida Gwaii & AK | Kyuquot/Quatsino | 61,882 | (32,052-85,984) |
| M | B | Haida Gwaii & AK | Juan de Fuca | 61,882 | (44,535-85,984) |
| M | B | Barkley/Clayoquot | BC North Coast | 61,882 | (61,882-85,984) |
| M | B | BC North Coast | Juan de Fuca | 53,209 | (44,535-61,882) |
| M | B | Barkley/Clayoquot | BC Central Coast | 44,535 | (23,067-61,882) |
| M | B | Kyuquot/Quatsino | BC North Coast | 44,535 | (44,535-61,882) |
| M | B | BC Central Coast | Juan de Fuca | 32,052 | (23,067-32,052) |
| M | B | Broughton Arch. | Barkley/Clayoquot | 23,067 | (11,947-32,052) |
| M | B | Broughton Arch. | Juan de Fuca | 19,834 | (16,601-23,067) |
| M | B | BC Central Coast | Kyuquot/Quatsino | 19,834 | (8,598-32,052) |
| M | B | BC Central Coast | BC North Coast | 14,274 | (6,188-23,067) |
| M | B | Haida Gwaii & AK | Broughton Arch. | 11,947 | (8,598-32,052) |
| M | B | Broughton Arch. | Kyuquot/Quatsino | 11,947 | (8,598-11,947) |
| M | B | Broughton Arch. | BC North Coast | 11,947 | (11,947-16,601) |
| M | B | Haida Gwaii & AK | BC Central Coast | 10,273 | (6,188-32,052) |
| M | B | Broughton Arch. | BC Central Coast | 8,598 | (6,188-11,947) |
| M | B | Haida Gwaii & AK | BC North Coast | 6,188 | (4,454-11,947) |
| M | B | Barkley/Clayoquot | Juan de Fuca | 4,454 | (3,205-6,188) |
| M | B | Kyuquot/Quatsino | Juan de Fuca | 4,454 | (4,454-11,947) |
| M | B | Barkley/Clayoquot | Kyuquot/Quatsino | 3,205 | (1,660-11,947) |
| M | W | Haida Gwaii & AK | Haida Gwaii & AK | 4,454 | (3,205-8,598) |
| M | W | BC North Coast | BC North Coast | 4,454 | (4,454-4,454) |
| M | W | BC Central Coast | BC Central Coast | 3,205 | (3,205-3,205) |
| M | W | Juan de Fuca | Juan de Fuca | 3,205 | (3,205-3,205) |
| M | W | Kyuquot/Quatsino | Kyuquot/Quatsino | 2,756 | (1,195-6,188) |
| M | W | Barkley/Clayoquot | Barkley/Clayoquot | 2,307 | (0-4,454) |

**Methods S1** Methodological details

*Sequencing of California Macrocystis*

For new *Macrocystis* samples from California (Supporting Information Table S1), blade samples were collected on scuba, by kayak, or intertidally depending on the location. To avoid collecting the same plant twice when sampling from the surface, we sampled a minimum of 5 m apart. We shipped silica-dried samples to the University of California, Davis, where we extracted DNA using the *Omega Bio-tek* E.Z.N.A. Mollusc DNA Kit (Cat. No. D3373) with the following modifications: 2% polyvinylpolypyrrolidone (PVPP) was added to the ML-1 Lysis buffer along with 0.1% 2-mercaptoethanol. Following digestion, the samples were spun, and the liquid was transferred to a clean tube before addition of C:I. Following extraction, we further purified samples using the *Qiagen* DNeasy PowerClean Cleanup Kit (Cat. No. 12877). We prepared libraries for whole-genome sequencing following the protocol of Rowan *et al.* (2019). Paired-end 150-bp libraries were sequenced on an *Illumina* NovaSeq X at the UC Davis Genome Center.

*Relationships within North American Macrocystis*

We constructed a phylogeny using *IQ-TREE* v.2.3.6 (Minh *et al.*, 2020). We began with the global 8x dataset and retained only the single individual with the highest depth per population. We then retained phylogenetically informative SNPs with a minimum count of three occurrences of the minor allele (*bcftools view --min-ac 3:minor*) and thinned to a minimum distance of 10-kbp between SNPs. We ran *IQ-TREE* using automated selection of the substitution model (Kalyaanamoorthy *et al.*, 2017) with correction for ascertainment bias (Lewis, 2001) (*-m MFP+ASC*), assessing branch support with 1000 replicates of the ultrafast bootstrap approximation (Hoang *et al.*, 2018) and SH-like approximate likelihood ratio tests (SH-aLRT; Guindon *et al.*, 2010) (*-B 1000 -alrt 1000*).

We tested for the possibility of gene flow between *Macrocystis* from different geographic regions of North America using *Dsuite* v.0.5 r53 (Malinsky *et al.*, 2021). *Dsuite* infers gene flow by calculating four-taxon Patterson's *D*-statistics (i.e., ABBA-BABA tests; Durand *et al.*, 2011) to detect excess allele sharing between pairs of taxa. We used the 8x *Macrocystis* dataset filtered to retain all California populations, the highest-diversity population from each group of geographically adjacent populations in AKBCWA, and a single Southern Hemisphere population (MP-CL-10) as the outgroup, requiring a minimum count of three occurrences of the minor allele and thinning to a minimum distance of 10 kbp between SNPs. We ran the *Dsuite Dtrios* command with default parameters using the phylogeny generated in *IQ-TREE* as the guiding topology, and then the *Dsuite Fbranch* command to infer specific branches at which gene flow occurred and to visualize results.

*Ecological niche modelling details*

We constructed ecological niche models (ENMs) using *MaxNet* v.0.1.4 (Phillips *et al.*, 2017) within the *R* package *SDMtune* v.1.3.1 (Vignali *et al.*, 2020). *MaxNet* is an *R* implementation of the original machine learning algorithm *Maxent* (Phillips *et al.*, 2006). We downloaded global occurrence records from the Global Biodiversity Information Facility (www.gbif.org) for *Nereocystis* and *Macrocystis*, as well as all brown algae (*Phaeophyceae*) occurrences in the Northeast Pacific (13.5º to 72.5º latitude and -180º to -91º longitude) to serve as background points (GBIF.org, 2024a,b,c). The use of brown kelp as background points was intended to reduce the effects of sampling bias in the occurrence records, assuming that the sampling bias in the occurrence records approximates the sampling bias in the background points.

We used a custom *R* script to filter occurrence records and background points. We first filtered *Nereocystis* and *Macrocystis* occurrences to the same geographic extent as the brown algae records. For all three taxonomic sets, we then retained only records corresponding to preserved specimens, removed records flagged with severe issues (invalid coordinates or geodetic datum; mismatches between coordinates, continent, and country; failed or suspicious coordinate reprojection; zero coordinates; and no taxon match or a match to a higher rank), and removed low-precision records inferred to have been originally recorded to <2 decimal places or to the nearest 10 or 15 arcminutes. We then individually removed any records outside the known range of *Nereocystis* or *Macrocystis*, any brown algae background records from areas biogeographically inaccessible to *Nereocystis* and *Macrocystis* (i.e., the Canadian Arctic and Atlantic Ocean), and any records located on land, thus retaining a total of 647 *Nereocystis*, 1,245 *Macrocystis*, and 43,782 brown algae occurrences.

We used environmental predictor variables from the MARSPEC dataset (Braconnot *et al.*, 2007; Sbrocco & Barber, 2013; Sbrocco, 2014) at 5-arcminute resolution. MARSPEC consists of bathymetric features plus bioclimatic variables related to sea surface temperature and salinity. We reprojected modern and LGM data (21 kya) at 5-arcminute resolution (equivalent to ~9.3 x 6.0 km at 50º latitude) to an Albers equal-area projection. For the LGM, we considered two different climatic datasets: the CCMS3 model and the LGM ensemble of all climate models except CCSM3. The CCSM3 model is the only model available in MARSPEC that accounts for increased salinity due to a drop in sea level during the LGM (Sbrocco, 2014), and we therefore evaluated the CCSM3 model separately from the remaining ensemble of models that failed to take this salinity effect into account.

In preparation for selecting environmental predictor variables and constructing ENMs, we created 20 resampled datasets of occurrences and background points, each time selecting a random combination of 80% of unique occurrences (i.e., occurrences occupying distinct raster cells in the environmental data; n = 198 and 226 prior to down-sampling for *Nereocystis* and *Macrocystis*, respectively) and 80% of unique background points (n = 1,540). To select a set of uncorrelated environmental predictor variables, we calculated the Spearman correlation coefficient ($\rho$) between variables in the modern data, excluding open ocean areas >12 km away from shore. We iteratively retained the single variable within each cluster of correlated variables ($\rho > 0.8$) that we arbitrarily judged to have the clearest conceptual link to habitat suitability for kelp, removing all other correlated variables within that cluster. From this set of uncorrelated variables, we then ran 20 initial *MaxNet* models in *SDMtune*, each time using one of the 20 different resampled datasets of occurrences and background points, and removed any variables with mean permutation importance below 5%.

After selecting the final environmental predictor variables, we turned hyperparameters for each species using the *gridSearch()* function in *SDMtune* with 5-fold cross-validation, and selected the model with the highest area under the receiver operating characteristic curve (AUC) in the test data. For both species, this procedure selected linear, quadratic, and product feature classes and regularization multiplier 0.2.

To run the final models for each species, we created 20 new resampled datasets of 80% occurrences and 80% background points as described above, and ran the *MaxNet* model in *SDMtune* 20 times using 5-fold cross-validation, with our final set of environmental predictor variables and hyperparameters. We assessed model performance using AUC and true skill statistic (TSS), and variable importance using permutation importance and a Jackknife test implemented in *SDMtune*. We projected each model to the modern, LGM CCSM3, and LGM ensemble (excluding CCSM3) time periods, and plotted the mean of the 20 models. We estimated a presence-absence threshold for each species as the habitat-suitability value in the current time period that would retain 95% of unique occurrence records.

*Pairwise directionality index details*

We calculated the directionality index ($\psi$; Peter & Slatkin, 2013) in AKBCWA using the 8x dataset for each species, filtered to minimum MAF of 0.05 and thinned to minimum 10 kbp between SNPs. We polarized SNPs based on inference of ancestral and derived alleles following Bemmels *et al.* (2025). The polarization method has previously been described in full (Bemmels *et al.*, 2025). Briefly, polarization involved aligning three outgroup genomes (split into 500-bp fragments) to the focal species' reference genome (outgroups: *Saccharina japonica* str. Ja from *Phycocosm* (Grigoriev *et al.*, 2021); *Laminara digitata* LdigPH10_18mv male genome v1.0 from *Phaeoexplorer* (Phaeoexplorer Project, 2024); and either *Macrocystis* or *Nereocystis* depending on the focal species) using *bwa-mem* v.0.7.17-r1188 (Li, 2013); filtering alignments for quality using *SAMtools* v.1.17 (Danecek *et al.*, 2021); converting to multi-species fasta format with *HTSBox* v.r345 (Li, 2012); and printing the alignments for the three outgroups using the *gerpcol* script v.2023/11/20 from Taylor et al. (2024). Then, we used a custom *R* script to define an allele as derived if it was not present in any of the three outgroups. If neither or both alleles met this definition or if no outgroup sequences aligned to the focal SNP, the derived allele could not be defined.

We considered using the time difference of arrival (TDoA) method (Peter & Slatkin, 2015) to statistically infer the geographic origin of range expansion, but reasoned that it is not well-suited to our study system for two main reasons. Firstly, the TDoA approach relies on correlations between geographic distance and $\psi$, but in the complex coastline of AKBCWA it is unlikely that distance between geographic coordinates as inferred from latitude and longitude in the *R* package *rangeExpansion* (Peter & Slatkin, 2015) would accurately reflect the minimum migration distance across the ocean between populations. This issue could be mitigated by customizing the standard approach to use a minimum ocean distance instead of geographic distance, but a second, more fundamental concern is that range expansions are not the only process than can cause non-zero $\psi$ values (Kemppainen *et al.*, 2024). While the impact of boundary effects has been explored (Kemppainen *et al.*, 2024), we reasoned that severe local population bottlenecks reflecting reductions in population size unrelated to geographic migration could also create false signals of range expansion by mimicking founder effects. This intuition was supported by preliminary results that revealed extreme $\psi$ values for comparisons involving bottlenecked populations. Contemporary population size, genetic diversity, and inbreeding rate estimates of *Nereocystis* and *Macrocystis* vary widely and suggest frequent population-specific bottlenecks (Bemmels *et al.*, 2025), which could distort $\psi$ values involving those populations and interfere with the TDoA approach.

Given these concerns, we opted not to use the formal TDoA method and instead to qualitatively infer expansion origins by plotting pairwise $\psi$ values on a map, as has been suggested for visualizing complex expansion scenarios (Peter & Slatkin, 2013). We calculated pairwise $\psi$ in *R* using the *get.all.psi.mc.bin()* function from Kemppainen *et al.* (2024). To reduce the effects of bottlenecks unrelated to range expansion, we reasoned that when geographically proximate populations show extremely different mean $\psi$ values (means across all possible pairwise comparisons), these differences are more likely due to recent bottlenecks experienced *in situ* rather than the founder effects of postglacial range expansion. We therefore grouped geographically proximate populations *ad hoc* and retained only the population 1 from each group with the lowest mean $\psi_{1,2}$ (where negative values of $\psi_{1,2}$ suggest migration from 1 to 2; Peter & Slatkin, 2013), thus retaining the population with the least evidence of post-migration bottlenecks from each group. We then visualized $\psi$ values between the retained populations on a map, for ease of interpretation plotting only $\psi$ between retained populations from geographically adjacent regions along the coastline.

*Ancestral recombination graph details*

We constructed ancestral recombination graphs (ARGs; Lewanski *et al.*, 2024) using *Relate* v.1.2.3 (Speidel *et al.*, 2019, 2021). We began with 8x datasets filtered to AKBCWA and removed SNPs for which the ancestral allele could not be determined (as described above in the *Pairwise directionality index details* section). We next retained first SNPs and then individuals with ≥90% non-missing data using *bcftools*. We phased and imputed missing data using *Beagle* v.5.5 (Browning *et al.*, 2018, 2021) with default parameters. We then converted VCF files to hap/sample format using *bcftools convert* and polarized SNPs using a custom python script and the previously described ancestral allele definitions.

As *Relate* requires a recombination map, we inferred one for each species using *FastEPRR* v.2.0 (Gao *et al.*, 2016). We reused a recombination map inferred from a previously published SNP dataset for BC and Washington (Bemmels *et al.*, 2025), without any missing data filter applied so that the genome would not be sporadically missing true SNP sites. We reduced datasets to one non-selfed individual per population for *Nereocystis* (n = 68) and up to two non-selfed individuals (if available) for *Macrocystis* (n = 67), respectively, retaining the individuals with the least amount of missing data. We removed singletons (ignored by *FastEPRR*), and then imputed and phased missing genotypes using *Beagle*. We ran *FastEPRR* using window and step sizes of 50 and 25 kbp, respectively, two custom training sets ("0;1;2;5;10;20;50;100;200" and "210;250;300;400;600;1000"), and re-estimation of variable recombination rates within each window. To convert recombination rates from $\rho$ into units of cM/Mb in *FastEPRR*, we assumed an effective population size ($N_e$) of $10^5$ for *Nereocystis* and $10^4$ for *Macrocystis*, which were rough estimates approximately an order of magnitude greater than previous $N_e$ estimates for individual populations (Bemmels *et al.*, 2025). Importantly, although the true $N_e$ is not known, the absolute magnitudes of recombination rate (which depend on $N_e$) are likely to have little impact on downstream analyses as they are multiplied by an arbitrary constant *R* in *Relate* to obtain transition probabilities in the Hidden Markov Model, and inferred tree topologies are robust to a wide range of *R* values (Speidel *et al.*, 2019). We converted *FastEPRR* output into the format required by *Relate* using a custom *R* script, and smoothed over any windows with a recombination rate of zero by converting any such windows and the adjacent two windows bounding it on either side to the mean recombination rate across all three windows.

To create a genomic mask file for *Relate* specifying which regions should be excluded due to not passing filtering criteria, we created a list of invariant sites filtered the same way as our variant sites (except that filters applicable only to variant sites were not applied). This list included sites initially identified as invariant across all individuals, as well as sites that became invariant after removing individuals that did not pass the filter requiring ≥90% non-missing data. All genomic positions that were not included in the filtered list of variant or invariant sites were masked. We then created a *.dist* file specifying the distance between SNPs after adjusting for masking using the *Relate* function *RelateFileFormats --mode FilterHapsUsingMask*.

We constructed an initial ARG for each chromosome using the function *Relate --mode all*, with a prior on the mutation rate of $8.135 \times 10^{-10}$ for both species, following Bemmels *et al.* (2025), and a prior on haploid effective population size of 2N = 200,000 and 20,000 for *Nereocystis* and *Macrocystis*, respectively. We then simultaneously re-estimated branch lengths and estimated effective population sizes ($N_e$) and mutation rates using the *Relate* script *EstimatePopulationSize.sh*. We excluded selfed individuals from each population because frequent selfing in both species (Bemmels *et al.*, 2025) is likely to reflect non-random mating as most zoospores settle within a few metres of their parent (Gaylord *et al.*, 2006; Edwards, 2022). Inclusion of selfed individuals produced by non-random mating could therefore bias estimates of coalescence rates and thus population sizes through time (Mather *et al.*, 2020). Given the much shorter generation time for kelp than for humans, we used custom time bins spanning $10^2$ to $10^6$ years before present (*--bins 2,6,0.142857142*). We used a generation time of one year for *Nereocystis* as it is an annual species. The average generation time is unknown in *Macrocystis*, which may live up to seven years (Dayton *et al.*, 1984). Most *Macrocystis* individuals do not survive beyond one year (Dayton *et al.*, 1984) but established individuals may shade out juvenile competitors (Bell & Siegel, 2022) and likely have a higher reproductive output given that reproductive output is strongly linked to biomass (Reed, 1987). We therefore chose a generation time of two years in *Macrocystis*, but caution that the rescaling from generations to absolute number of years should be considered approximate in this species.

We additionally repeated the above ARG-reconstruction procedure and analyses for a global dataset of *Macrocystis*, to permit estimation of divergence times between global regions. We began with the global 8x dataset and subsetted it to include all Chilean and Californian populations plus one population per genetic cluster from AKBCWA (populations MP-HG-01, -NC-01, -CC-01, -BA-07, -QS-01, -CS-04, and -WC-01), resulting in a globally balanced distribution of samples (58 and 61 individuals from the Southern and Northern Hemispheres, respectively). We then followed the same methods as for the AKBCWA dataset except that we did not remove individuals with <90% non-missing data, as unequal sequencing depth would have resulted in removing some entire populations from California.
